# Mutations in *NAKED-ENDOSPERM* IDD genes reveal functional interactions with *SCARECROW* and a maternal influence on leaf patterning in C_4_ grasses

**DOI:** 10.1101/2022.10.28.514209

**Authors:** Thomas E. Hughes, Olga Sedelnikova, Mimi Thomas, Jane A. Langdale

## Abstract

Leaves comprise a number of different cell-types that are patterned in the context of either the epidermal or inner cell layers. In grass leaves, two distinct anatomies develop in the inner leaf tissues depending on whether the leaf carries out C_3_ or C_4_ photosynthesis. In both cases a series of parallel veins develops that extends from the leaf base to the tip but in ancestral C_3_ species veins are separated by a greater number of intervening mesophyll cells than in derived C_4_ species. We have previously demonstrated that the GRAS transcription factor SCARECROW (SCR) regulates the number of photosynthetic mesophyll cells that form between veins in the leaves of the C_4_ species maize, whereas it regulates the formation of stomata in the epidermal leaf layer in the C_3_ species rice. Here we show that SCR is required for inner leaf patterning in the C_4_ species *Setaria viridis* but in this species the presumed ancestral stomatal patterning role is also retained. Through a comparative mutant analysis between maize, setaria and rice we further demonstrate that loss of NAKED-ENDOSPERM (NKD) INDETERMINATE DOMAIN (IDD) protein function exacerbates loss of function *scr* phenotypes in the inner leaf tissues of maize and setaria but not rice. Specifically, in both setaria and maize, *scr;nkd* mutants exhibit an increased proportion of fused veins with no intervening mesophyll cells, whereas inner leaf tissues are patterned normally in *scr;nkd* mutants of rice. Thus, combined action of SCR and NKD may control how many mesophyll cells are specified between veins in the leaves of C_4_ but not C_3_ grasses. Finally, we identified a maternal effect in maize in which maternally derived NKD can affect patterning of cells in leaf primordia that are initiated during embryogenesis. Together our results provide insight into the evolution of cell patterning in grass leaves, demonstrate a novel patterning role for IDD genes in C_4_ leaves and suggest that NKD can influence embryonic leaf development non-cell autonomously from the surrounding maternal tissue.

**Summary statement:** Mutations in *NKD* IDD genes enhance loss of function *scr* phenotypes in the leaves of C_4_ grasses maize and *Setaria viridis* but not in the C_3_ grass rice, and reveal a maternal effect on cell-type patterning in leaves that are initiated during embryogenesis.

## INTRODUCTION

Understanding how cell patterning is genetically regulated is a key challenge in developmental biology. In grass leaves, two distinct cellular anatomies underpin photosynthesis. In grasses such as rice (*Oryza sativa*) that carry out C_3_ photosynthesis, widely spaced parallel veins are encircled by non-photosynthetic bundle-sheath cells which are themselves separated by up to ten photosynthetic mesophyll cells. By contrast, in grasses that perform C_4_ photosynthesis, such as maize (*Zea mays*) and green millet (*Setaria viridis*), parallel veins are surrounded by concentric layers of bundle-sheath and mesophyll cells, both of which are photosynthetic. This arrangement of cell-types is referred to as ‘Kranz’ because the two cell-types form wreaths around the veins and Kranz is German for wreath (Haberlandt, 1896). Notably Kranz anatomy evolved from C_3_-type anatomy on multiple independent occasions (Grass Phylogeny Working Group II, 2012; Sage et al., 2011), each occurrence generating leaves with higher vein densities than in the ancestral form because bundle-sheath cells are separated by fewer mesophyll cells. To date, very few regulators of cell-patterning in inner leaf tissues have been identified in C_3_ or C_4_ grass species.

We previously demonstrated that duplicate genes encoding the GRAS transcription factor SCARECROW (SCR) regulate cell divisions in the innermost ground meristem layer of maize leaf primordia to determine the number of M cells that form between veins (Hughes et al., 2019). In double *Zmscr1;Zmscr1h* mutants, the majority of bundle-sheath cells are separated by one rather than two mesophyll cells, and in some cases bundle-sheath cells are fused with no intervening mesophyll cells. In addition, many veins develop ectopic sclerenchyma either ad- or abaxially and some veins are surrounded by additional bundle-sheath cells that are not in contact with the vasculature. Intriguingly, no such patterning perturbations were observed in inner leaf tissues when *SCR* orthologs were mutated in rice (Hughes & Langdale, 2022). Instead, loss of function mutants in rice fail to develop stomata on the leaf surface (Hughes & Langdale, 2022; Wu et al., 2019). The distinction between mutant phenotypes in maize and rice leaves raises the possibility that the deployment of SCR function in inner leaf tissues was associated with the evolution of Kranz anatomy, with the patterning of stomata in the epidermis being the ancestral role in grass leaves.

Although the function of SCR itself could differ between maize and rice, for example by targeting different downstream genes, the distinct patterning roles observed could alternatively result from species-specific differences in SCR interacting proteins. Little is known about how SCR-mediated patterning is regulated in monocots but in Arabidopsis roots, SCR functions with another GRAS transcription factor (SHORTROOT (SHR)) and with several INDETERMINATE DOMAIN (IDD) C2H2 zinc-finger transcription factors (Laurenzio et al., 1996; Long et al., 2015; Nakajima et al., 2001; Ogasawara et al., 2011; Welch et al., 2007). The IDD genes (also referred to as BIRD genes), act both to modulate *SCR* and *SHR* gene expression and to co-regulate the expression of downstream target genes (Welch et al., 2007). Despite the existence of many IDD genes in grass genomes, and the demonstration that *SCR* and *SHR* have patterning functions in maize and rice, links between *SCR* and IDD genes in the patterning of either root or leaf cell-types in monocots have not yet been established.

We previously proposed that the *NAKED-ENDOSPERM* (*NKD*) *IDD* genes (originally referred to as *ZmJAY* genes) may function during Kranz patterning in maize (Fouracre et al., 2014; Sedelnikova et al., 2018). This proposal was based on the fact that *ZmNKD1* and *ZmNKD2* transcripts accumulate during early leaf development at the relevant stages of Kranz patterning (Wang et al., 2013), both transcripts accumulate specifically in mesophylls of mature leaves (Chang et al., 2012; Li et al., 2010) and both have known patterning functions in aleurone development (Gontarek et al., 2016; Yi et al., 2015). Building on this suggestion and on the outstanding questions raised above, here we have tested two hypotheses. First, that SCR function is required for patterning inner leaf tissues in C_4_ grasses and second that NKD is a component of the patterning pathway. By characterizing *scr* mutants in the C_4_ species *Setaria viridis* and carrying out a comparative analysis of *scr;nkd* mutants in maize, *Setaria viridis* and rice, we found evidence to support both hypotheses. In addition, we identified a maternal effect of NKD on the patterning of leaves formed during embryogenesis in maize.

## RESULTS

### SCR patterns both epidermal and inner tissues in leaves of Setaria viridis

To determine whether the distinct patterning roles of SCR in the rice epidermis versus the maize inner leaf reflect C_3_-versus C_4_-specific functions, we generated loss-of-function mutants in the C_4_ species *Setaria viridis* (hereafter referred to as setaria) using a CRISPR/Cas9 system (Fig. 1A, Fig. S1). As in maize and rice, two *SCR* genes are present in setaria (*SvSCR1*-Sevir.7G316501 and *SvSCR2*-Sevir.8G008100), each pair the result of a within species duplication (Hughes et al., 2019). In gene edited T2 lines, a proportion of plants were identified that were extremely stunted, paler than wild-type and did not survive beyond 3-4 weeks after germination (Fig. 1B-D). These plants appeared at the expected segregation ratios of 1/4 or 1/16 depending on whether the parental plant was *Svscr1/+*;*Svscr2* (or *Svscr1*;*Svscr2/+*) or *Svscr1/+*;*Svscr2/+* respectively. In all cases, these phenotypically abnormal plants were confirmed to be homozygous for both *Svscr1* and *Svscr2*, with phenotypically wild-type plants always being heterozygous or wild-type for one of the *SCR* genes. Because single *Svscr* mutants displayed no growth perturbations (Fig. S2) and single mutants in both rice and maize were not associated with any patterning defects, all further analyses were undertaken with homozygous double mutants. The perturbed growth phenotype exhibited by *Svscr1;Svscr2* plants was far more severe than that seen in maize *Zmscr1;Zmscr1h* mutants (where plants can be grown for 6-8 weeks in the greenhouse without issue) (Hughes et al., 2019), but was similar to the phenotype observed in *Osscr1*;*Osscr2* mutants of rice (Hughes & Langdale, 2022). We therefore reasoned that, as in rice, *SCR* may play a role in stomatal patterning in setaria. Epidermal impressions of *Svscr1;Svscr2* mutant leaves confirmed this hypothesis, revealing an absence of stomata on both the abaxial and adaxial leaf surfaces (Fig. 2A-F). Therefore, the stomatal patterning role first identified in rice is not a C_3_-specific function.

**Figure 1.**
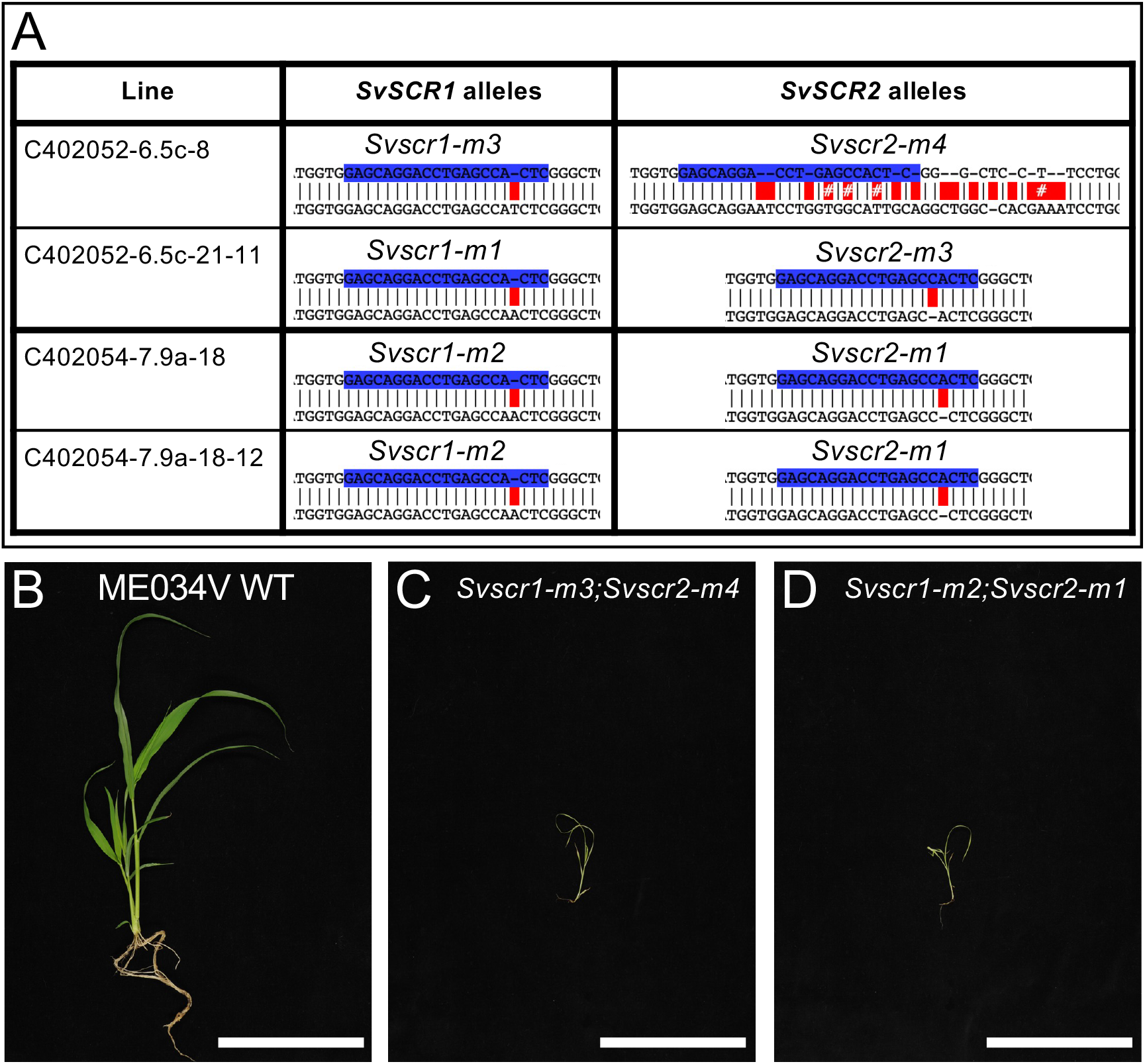
Mutations in *SvSCR1* and *SvSCR2* affect plant growth. **A)** Sequences of mutant *Svscr1* and *Svscr2* alleles in two independent lines (C_4_02052-6 & C_4_02054-7). Wild-type (WT) sequence is shown on top, with the sequence of the mutant allele shown beneath. Guide sequences are depicted in blue, and mismatches between the WT and mutant sequence indicated in red. **B-D)** Photos of WT ME034V (B), *Svscr1-m3;Svscr2-m4* (C) and *Svscr1-m2-Svscr2-m1* (D) plants taken 20 days after sowing. Scale bars: 10cm.

**Figure 2.**
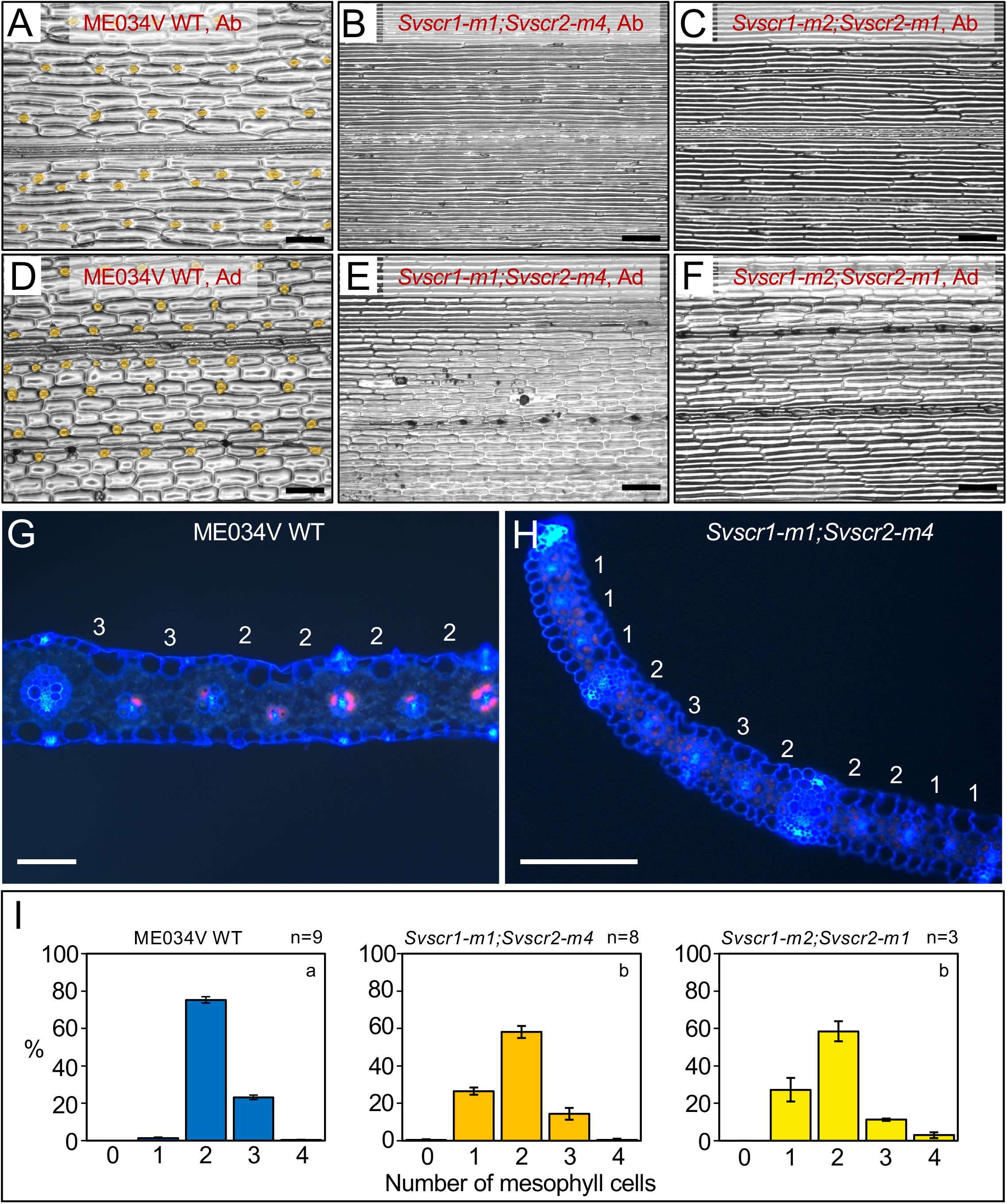
*Svscr1;Svscr2* mutant plants have no stomata and fewer mesophyll cells separating veins. **A-F)** Stomatal impressions of wild-type (WT) ME034V (A,D), *Svscr1-m1;Svscr2-m4* (B,E) and *Svscr1-m2;Svscr2-m1* (C,F) leaf 3 from either the abaxial (A-C) or adaxial (D-F) surface. Stomata are false coloured orange. Scale bars: 100μm. **G-H)** Transverse sections imaged using UV illumination of WT ME034V (G) and *Svscr1-m1;Svscr2-m4* (H) leaf 4, taken from the mid-point along the proximal-distal axis. The number of mesophyll cells between each pair of veins is indicated above the leaf. Scale bars: 100μm. **I)** Histograms summarizing the mean number of mesophyll cells separating veins in WT ME034V and two independent *Svscr1;Svscr2* mutants. The number of biological replicates is indicated above each plot and letters in the top right corner of each plot indicate statistically different groups (*P*≤0.05, one-way ANOVA and Tukey’s HSD) calculated using the mean number of mesophyll cells in each genotype.

To determine whether *SCR* also plays a role in inner leaf patterning in setaria, transverse sections of wild-type and *Svscr1;Svscr2* mutant leaves were examined (Fig. 2G, H). In ME034V wild-type leaves, around 70-80% of veins were separated by two mesophyll cells with the rest separated by three (a feature that is more common in setaria than in maize) (Fig. 2I). In contrast, 20-30% of veins in *Svscr1;Svscr2* mutant leaves were separated by just a single M cell (Fig. 2I). These data demonstrate that, as in maize, *SvSCR* genes regulate cell divisions in the ground meristem to determine how many mesophyll cells develop between bundle-sheath cells. Intriguingly, *SvSCR* genes undertake this inner leaf patterning role in addition to a role in stomatal patterning.

### A feedback loop links SCR and NKD IDD gene expression in maize

The distinct roles of *SCR* genes in maize, rice and setaria – patterning inner, epidermal or both tissue layers of the leaf respectively – could be due to species-specific differences in the activity of interacting IDD genes, of which *NKD* genes are potential candidates. To determine whether the expression of *ZmSCR1/h* genes is interconnected with *ZmNKD1/2* gene function in maize (as is the case for *SCR* and IDD genes in Arabidopsis), we first quantified transcript levels of each gene in existing double *Zmscr1-m2;Zmscr1h-m1* (Hughes et al., 2019) and double *Zmnkd1-Ds;Zmnkd2-Ds* (Yi et al., 2015) mutants (Fig. 3A, Fig. S3). *ZmSCR1* and *ZmSCR1h* transcripts accumulated at elevated levels in the *Zmnkd1-Ds;Zmnkd2-Ds* mutant, in both cases being increased around two-fold relative to wild-type (Fig. 3A). In contrast, *ZmNKD1* and *ZmNKD2* transcripts accumulated at lower levels in *Zmscr1-m2;Zmscr1h-m1* mutants than in wild-type, with *ZmNKD1* reduced on average to 60% and *ZmNKD2* to 50% of wild-type levels (Fig. 3A). To determine whether the interactions between NKD and SCR are likely to occur within or between cell-types, *in situ* hybridization was carried out to localize transcripts in developing leaf primordia. Figure 3B shows that both *NKD1* and *SCR1* transcripts accumulate in the ground meristem cells that surround developing vascular centres. Together these data suggest that a negative feedback loop controls *ZmSCR1/1h* and *ZmNKD1/2* transcript levels, most likely in a cell-type specific manner, with ZmSCR1/ZmSCR1h promoting the expression of *ZmNKD1*/*ZmNKD2*, and ZmNKD1/ZmNKD2 repressing the expression of *ZmSCR1*/*ZmSCR1h*.

**Figure 3.**
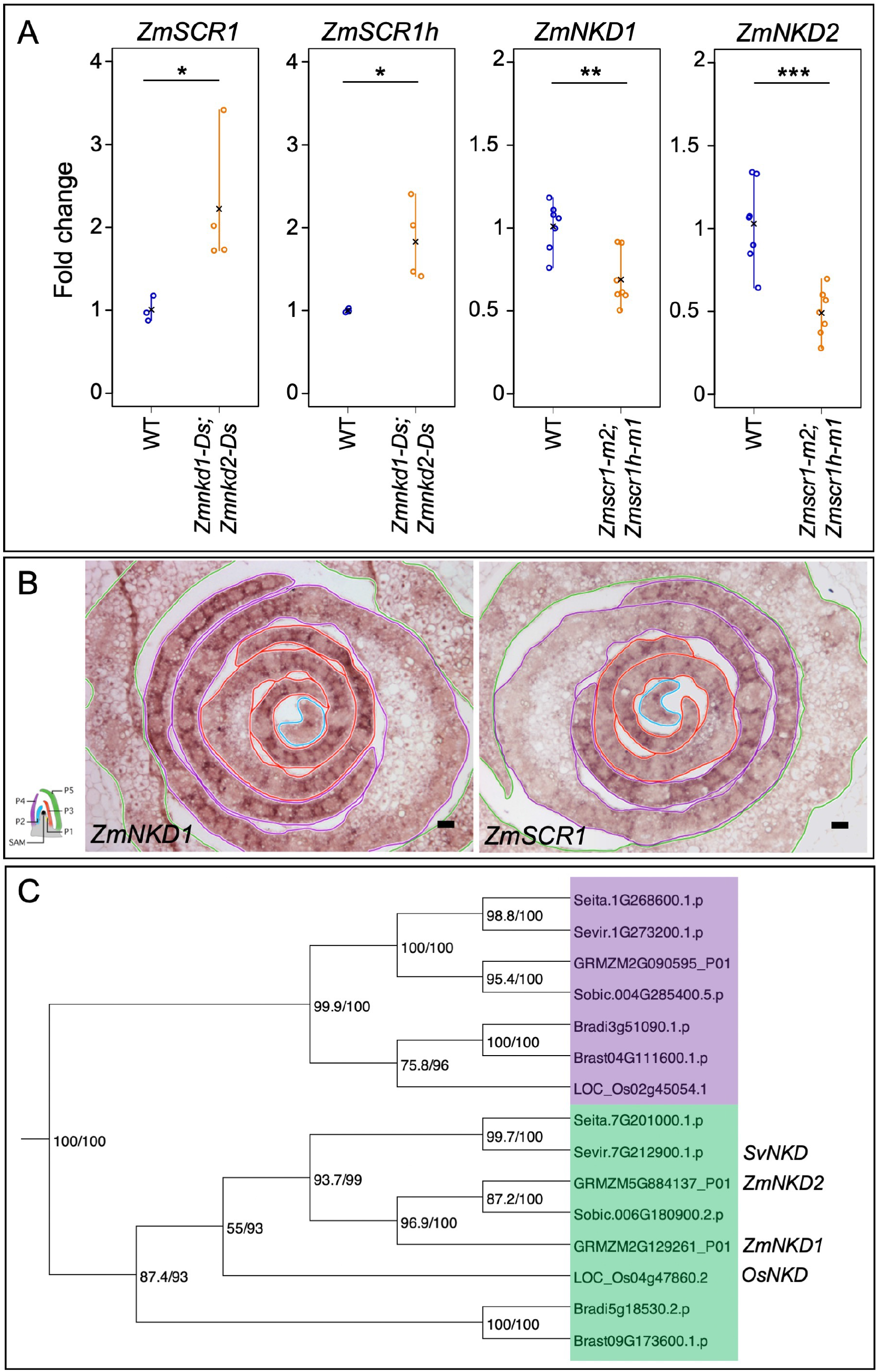
*NKD* and *SCR* transcripts accumulate in the same spatial domain with levels determined by a feedback loop. **A)** Quantitative RT-PCR of *ZmSCR1* and *ZmSCR2* transcripts in the *Zmnkd1-Ds;Zmnkd2-Ds* mutant, and *ZmNKD1* and *ZmNKD2* in the *Zmscr1-m2;Zmscr1h-m2* mutant. Open circles are individual biological replicates, black crosses indicate the mean for each genotype. Statistical significance as calculated on log-transformed fold change data by Student’s *t*-test (two-tailed) indicated above each plot: \**P*≤0.05; \*\**P*≤0.01; \*\*\**P*≤0.001. **B)** *In-situ* hybridization to *ZmNKD1* and *ZmSCR1* in maize wild-type B73 apices. In each image the P2 primordium is outlined in blue, P3 in red, P4 in purple and P5 in green, as indicated in the adjacent cartoon diagram. Darker purple signal represents successful hybridization to each transcript of interest. Scale bars: 50μm. **C)** Maximum likelihood phylogeny of the *NKD* genes in monocots. The *NKD* clade is highlighted in green, and the adjacent monocot clade in purple. Bootstrap values are displayed at each branch of the phylogeny.

### Loss of function nkd mutants do not exhibit perturbed leaf development

Given the patterning role of SCR in leaves of maize, setaria and rice, and the interaction between *SCR* and *NKD* gene expression in maize, we next sought to determine the role of *NKD* in leaf development in each of the three species. To this end, we first identified *NKD* orthologs in setaria and rice by constructing a maximum likelihood phylogeny. Figure 3C shows that *NKD* exists as a single copy gene in both setaria and rice (*SvNKD* – Sevir.7G212900, *OsNKD* – LOC_Os04g47860), suggesting that the two copies in maize (*ZmNKD1* and *ZmNKD2*) resulted from the recent whole genome duplication (Swigonova et al., 2004). For phenotypic characterization, the homozygous double *Zmnkd1-Ds;Zmnkd2-Ds* line and newly generated CRISPR/Cas9 loss-of-function mutants in setaria and rice were all compared to wild-type lines from the same genetic background (Fig. 4A, 4B, Fig. S1 & S4). Homozygous *Zmnkd1-Ds;Zmnkd2-Ds* seed exhibited the characteristic shrunken kernel phenotype that is caused by defective patterning of the aleurone layer but no altered seed phenotype was observed in *Svnkd* or *Osnkd* mutants (Fig. S5). Overall plant growth was normal in maize, setaria and rice *nkd* mutants (Fig. 4B) and no leaf patterning perturbations were observed (Fig. 4C-J, Fig. S6). In *Zmnkd1-Ds;Zmnkd2-Ds* mutants, no changes relative to wild-type were observed in leaf vein-density or in the ratio of rank-1 to rank-2 intermediate veins (Fig. 4G & 4H), traits that are both altered in *Zmscr1;Zmscr1h* mutants. Furthermore, there was no change in the number of mesophyll cells separating veins (the most penetrant phenotype in equivalent *scr* mutants) in either maize or setaria *nkd* mutants (Fig. 4I & 4J). Taken together, these results indicate that *NKD* is not necessary for normal leaf development in maize, setaria or rice.

**Figure 4.**
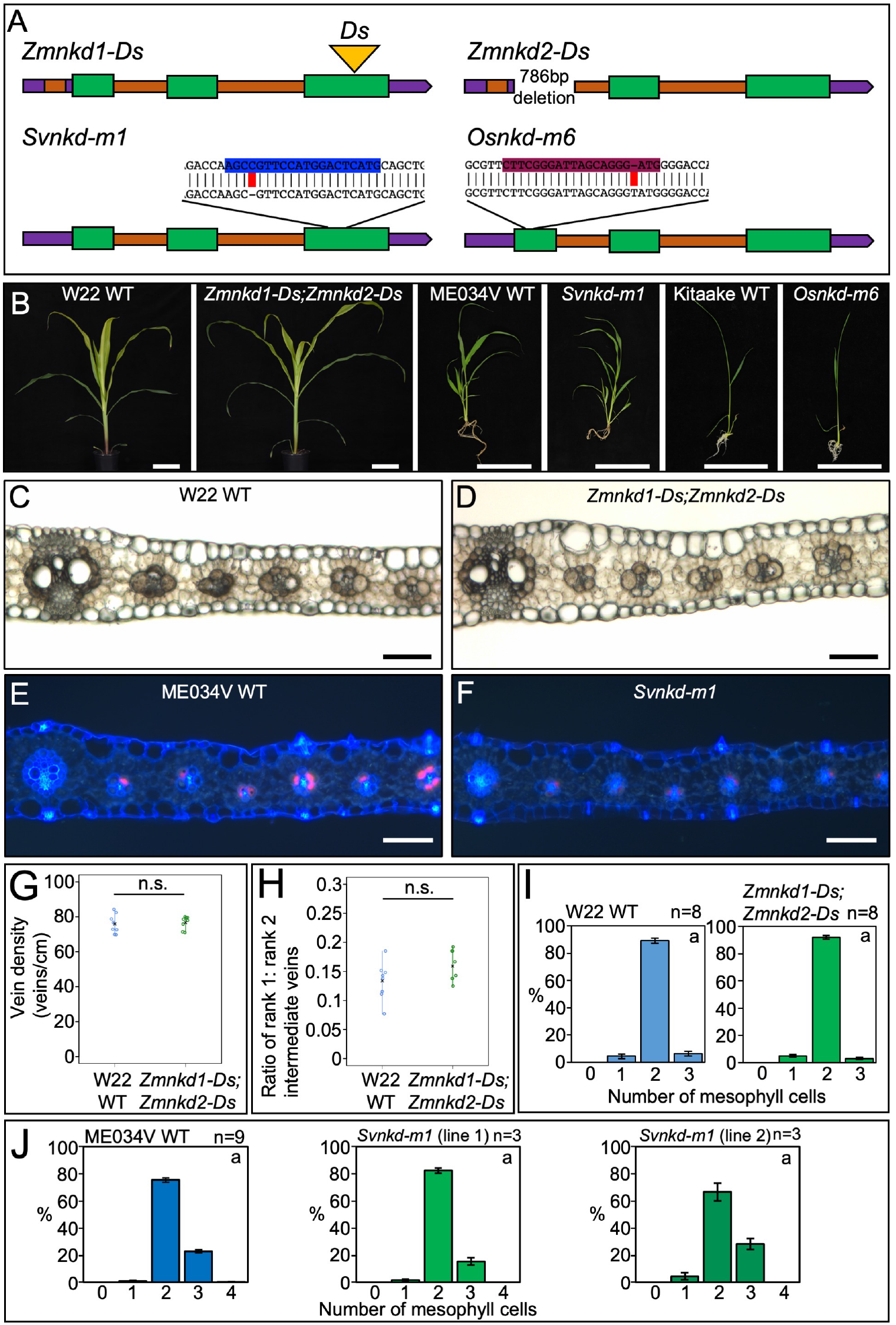
*nkd* loss of function mutations do not perturb growth in maize, rice or setaria. **A)** Cartoon depictions of loss of function *nkd* alleles in maize (*Zm*), setaria (*Sv*) and rice (*Os*). In each case, 5’ and 3’ untranslated regions are depicted in purple, introns in orange and coding regions in green. Transposon insertions are indicated by a yellow triangle. For setaria and rice, CRISPR/Cas9 guide sequences are indicated in blue and maroon respectively, with the sequence of the mutant allele beneath with edits highlighted in red. **B)** Whole plant phenotypes of maize, setaria and rice *nkd* mutants. Photos were taken 31 days (maize), 20 days (setaria) or 14 days (rice) after sowing. Scale bars: 10cm. **C-F)** Transverse sections of maize wild-type (WT) W22 (C), *Zmnkd1-Ds;Zmnkd2-Ds* (D), setaria WT ME034V (E) and *Svnkd-m1* (F), imaged under either brightfield (maize) or UV illumination (setaria). Images were taken at the mid-point along the proximal-distal axis of either leaf 5 (maize) or leaf 4 (setaria). Scale bars: 100μm. **G-H)** Quantification of vein density (G) and the ratio of rank1:rank2 intermediate veins (H) of WT W22 (blue) and *Zmnkd1-Ds;Zmnkd-Ds* (green) mutants. Open circles indicate distinct biological replicates, black crosses indicate the mean for each genotype. Statistical significance as calculated by Student’s *t*-test (two-tailed) indicated above each plot: n.s. *P*>0.05. **I-J)** Histograms summarising the mean number of mesophyll cells separating veins in WT W22 versus *Zmnkd1-Ds;Zmnkd2-Ds* (I) and WT ME034V versus two independent *Svnkd-m1* lines (J). Error bars are the standard error of the mean, and the number of biological replicates is indicated above each plot. Letters in the top right corner of each plot indicate statistically different groups (*P*≤0.05, one-way ANOVA and Tukey’s HSD) calculated using the mean number of mesophyll cells in each genotype. WT is coloured blue and *nkd* mutants green.

### Loss of NKD gene function enhances scr mutant phenotypes in maize and setaria but not in rice

To determine whether a role for *NKD* in leaf patterning could be revealed in the absence of SCR function, quadruple *scr1;scr1h;nkd1;nkd2* mutants of maize and triple *scr1;scr2;nkd* mutants of setaria and rice were generated (Fig. 5, Fig. S1 & S4). An initial assessment of overall growth phenotypes revealed that perturbations in three independent triple *Osscr1;Osscr2;Osnkd* mutant lines were similar to those seen in double *Osscr1;Osscr2* mutants (Fig. 5B-D). By contrast, triple *Svscr1;Svscr2;Svnkd* mutants displayed more severe perturbations than double *Svscr1;Svscr2* mutants (Fig. 5E-I). In the progeny of *Svscr1;Svscr2/+;Svnkd/+* plants from two independent lines, the resultant triple mutants were smaller and died faster than double *Svscr1;Svscr2* mutant siblings (Fig. 5H & 5I). This was consistent in lines fixed for *Svnkd*, where only 8/20 and 8/34 triple mutants from two independent *Svscr1;Svscr2;Svnkd* lines survived 22 days after sowing, compared to 14/15 and 13/17 *Svscr1;Svscr2* double mutants. In maize, quadruple *Zmscr1-m2;Zmscr1h-m1;Zmnkd1-Ds;Zmnkd2-Ds* mutants appeared similar to double *Zmscr1;Zmscr1h* mutants, with pale, drooping leaves and a shorter stature than either wild-type or double *Zmnkd1-Ds;Zmnkd2-Ds* mutant plants (Fig. 5J-O). However, germination of quadruple mutants was often inconsistent, and seedlings appeared weaker and more susceptible to fungal contamination. Mutant combinations thus demonstrate that NKD and SCR act synergistically in maize and setaria but not in rice.

**Figure 5.**
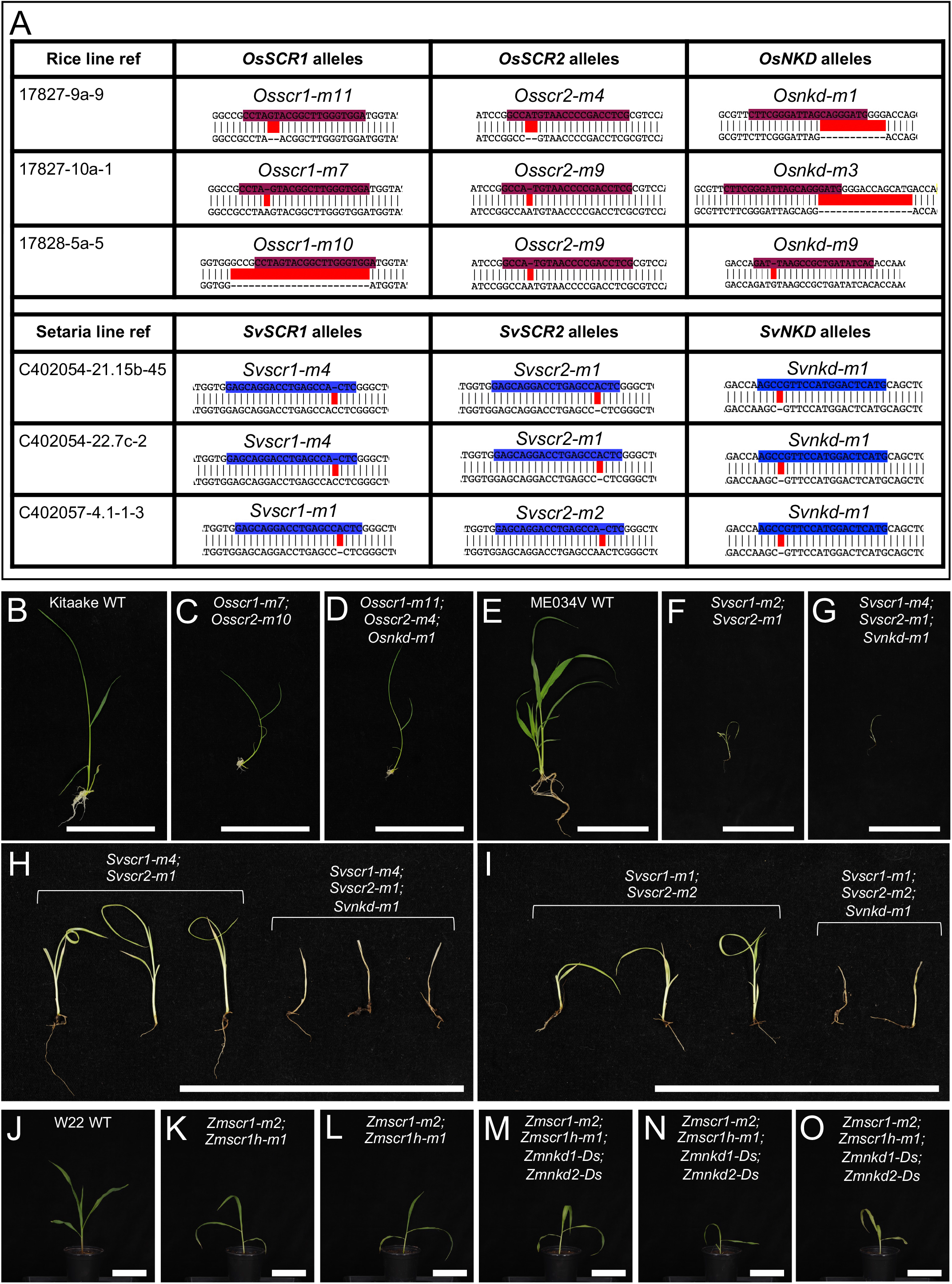
Loss of function *nkd* mutations enhance growth perturbations in *scr* mutants of maize and setaria but not rice. **A)** Sequences of mutant *scr1;scr2;nkd* alleles in three independent lines of both rice and setaria. Wild-type sequences are shown above with guides highlighted in either maroon (rice) or blue (setaria). Edits are highlighted in red. Note *Svscr1-m2* allele is included in Figure 1. **B-O)** Whole plant phenotypes of rice (B-D), setaria (E-I) and maize (J-O). Images were taken 14, 20 or 21 days after sowing for rice, setaria and maize respectively. Plants in (H) and (I) are from segregating families in each case, such that *Svscr1;Svscr2* and *Svscr1;Svscr2;Svnkd* in each panel are siblings. Scale bars: 10cm.

To determine the impact of SCR and NKD interactions on leaf development, inner leaf patterning was examined in quadruple (maize) and triple (setaria and rice) mutants. Notably, the complete penetrance of the stomatal phenotype in double *scr1;scr2* mutants of setaria and rice precluded an assessment of interactions with *NKD* during stomatal development but no changes in stomatal patterning were observed in single *Svnkd* or *Osnkd* mutants (Fig. S6). In inner leaf tissues, the number of mesophyll cells between veins were the same in three independent triple *Osscr1;Osscr2;Osnkd* mutant lines of rice as in corresponding double *Osscr1;Osscr2* mutants, with the modal number of mesophyll cells being five in all cases (Fig. 6A-C). By contrast, fused veins (where the bundle-sheath cells of adjacent veins are in contact with no intervening mesophyll cells) were observed in triple *Svscr1;Svscr2;Svnkd* mutants of setaria but not in double *Svscr1;Svscr2* mutants (Fig. 6D-F). Such fusions were not found in every triple mutant examined, with examples observed in 10/22 samples across two experiments with two independent lines (Fig 6F & Fig. S7). However, only one occurrence of a fused vein was observed in twenty-five double *Svscr1;Svscr2* mutant samples. Fused veins were also observed in quadruple mutants of maize (Fig. 7A-H). Quadruple mutants did not exhibit higher vein density or an increase in veins associated with sclerenchyma when compared to double *Zmscr1-m2;Zmscr1h-m1* mutants (Fig. 7I & 7J) but the increase in the number of veins fused to adjacent veins was consistent and statistically significant (Fig. 7F, 7H, 7K-M). Furthermore, whereas only two adjacent veins fused in double *Zmscr1-m2;Zmscr1h-m1* mutants, groups of three or more fused veins were often seen in quadruple mutants (Fig. 7L). These fused veins resulted in a large increase in the number of veins separated by no mesophyll cells, with only ~20% of veins separated by the normal two mesophyll cells in quadruple mutants, compared to just over 40% in double *Zmscr1-m2;Zmscr1h-m1* mutants. In both maize and setaria the proportion of veins separated by only one mesophyll cell was similar in double and quadruple/triple mutants (Fig. 6F & 7M). To ensure that these differences were not caused by heterozygosity at unlinked loci following the cross between the *Zmscr1-m2;Zmscr1h-*m1 and *Zmnkd1-Ds;Zmnkd2-D* double mutants, we also compared *Zmscr1-m2;Zmscr1h-m1* mutants pre- and post-outcross and found no difference in the proportion of fused veins (Fig. S8A). Furthermore, we validated that the phenotype was consistent across two further quadruple mutants from crosses using independent *Zmnkd1-Ds;Zmnkd2-Ds* plants (Fig. S8B). Together these results demonstrate that in the C_4_ species maize and setaria, but not in the C_3_ species rice, *NKD* genes function with *SCR* to determine whether, and how many, mesophyll cells are positioned between veins.

**Figure 6.**
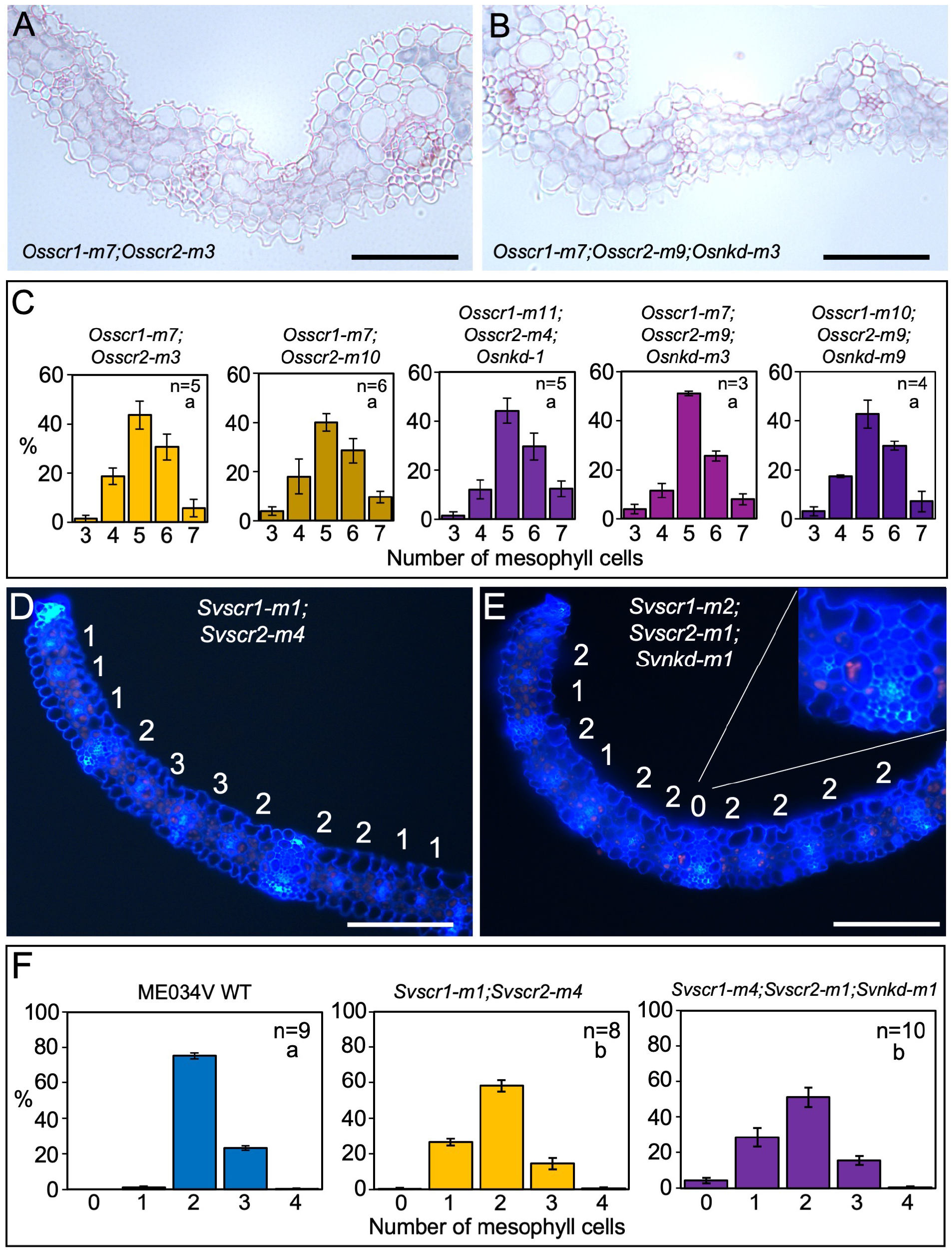
Loss of function *nkd* mutations induce the formation of fused leaf veins in *scr* mutants of setaria but not rice. **A-B)** Transverse sections of *Osscr1-m7;Osscr2-m3* (A) and *Osscr1-m7;Osscr2-m9;Osnkd-m3* (B) mutant leaves, taken at the midpoint along the proximal-distal axis of leaf 5. **C)** Histograms summarizing the mean number of mesophyll cells separating veins in two independent *Osscr1;Osscr2* (yellow) and three independent *Osscr1;Osscr2;Osnkd* (purple) mutant lines. Error bars are standard error of the mean. Sample sizes (n=) are biological replicates. **D-E)** Transverse sections of *Svscr1-m1;Svscr2-m4* (D) and *Svscr1-m2;Svscr2-m1;Svnkd-m1* (E) mutant leaves, taken at the midpoint along the proximal-distal axis of leaf 4, imaged under UV illumination. The number of mesophyll cells separating veins is displayed above each region. In (E) the inset shows an example of a fused vein with no separating mesophyll cells. **F)** Histograms summarizing the mean number of mesophyll cells separating veins in wild-type (WT) ME034V (blue), *Svscr1-m1;Svscr2-m4* (yellow) and *Svscr1-m4;Svscr2-m1;Svnkd-m1* (purple) mutants. Error bars are standard error of the mean. Sample sizes (n=) are biological replicates and letters in the top right corner (beneath sample sizes) of each plot indicate statistically different groups (*P*≤0.05, one-way ANOVA and Tukey’s HSD) calculated using the mean number of mesophyll cells in each genotype. Scale bars: 100 μm.

**Figure 7.**
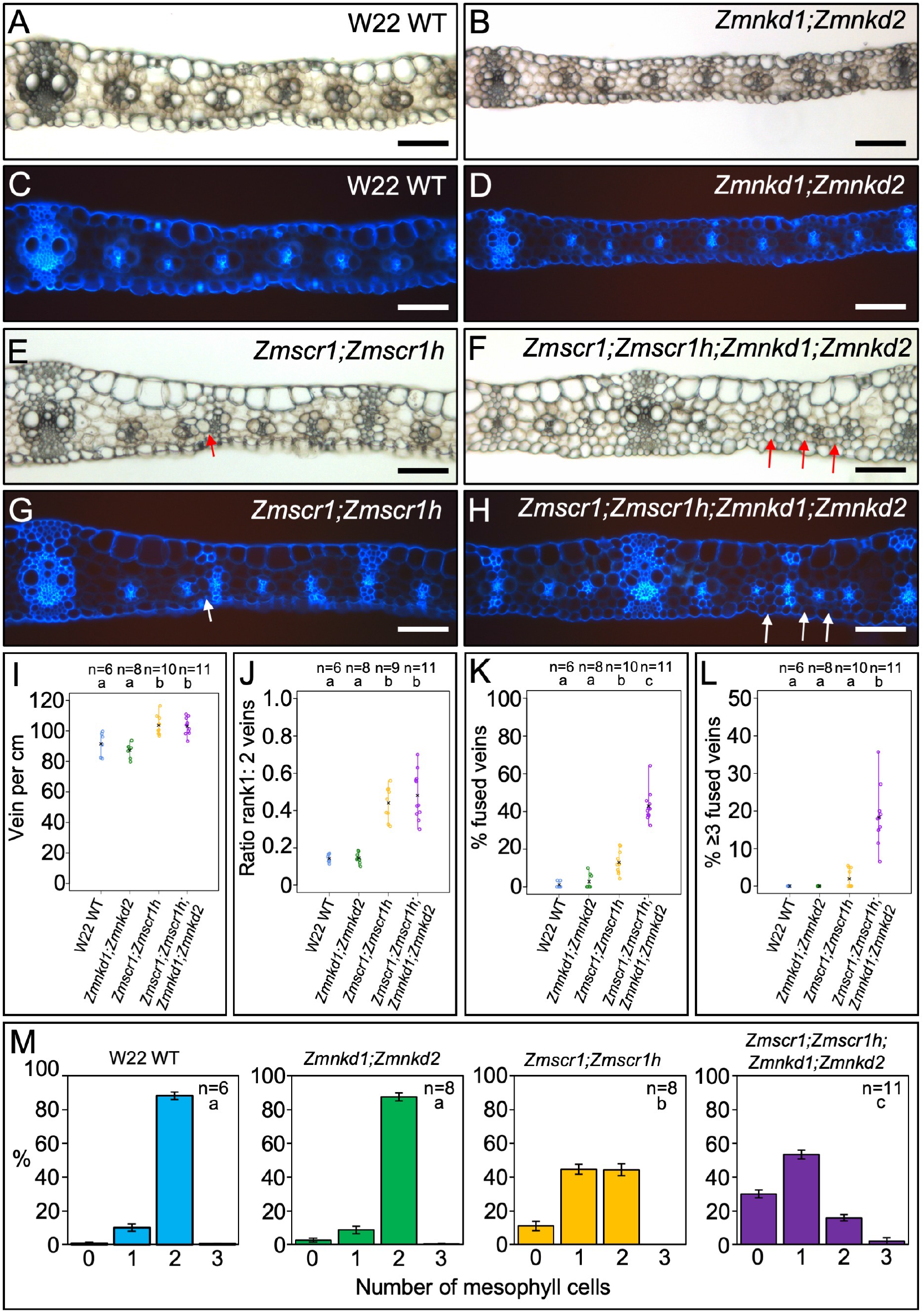
*Zmscr1;Zmscr1h;Zmnkd1;Zmnkd2* quadruple mutants have a striking increase in the number of fused leaf veins compared to *Zmscr1;Zmscr1h* double mutants. **A-H)** Transverse cross sections of wild-type (WT) W22 (A & C), *Zmnkd1-Ds;Zmnkd2-Ds* (B & D), *Zmscr1-m2;Zmscr1h-m1* (E & G) and *Zmscr1-m2;Zmscr1h-m1;Zmnkd1-Ds;Zmnkd2-Ds* (F & H) leaf 3, from the mid-point along the proximal distal axis and imaged under either brightfield (A-B, E-F) or UV (C-D, G-H) illumination. Arrows point to fused veins. Scale bars: 100 μm. **I-L)** Strip charts summarizing quantification of vein density (I), the ratio of rank 1: rank 2 intermediate veins (J), the % of fused veins (K) and the % of veins formed in runs of ≥3 fused veins (L). Open circles indicate measurements from independent biological replicates, and black crosses indicate the mean for each genotype. The number of biological replicates (n=) is indicated above each plot and letters at the top of each plot indicate statistically different groups (*P*≤0.05, one-way ANOVA and Tukey’s HSD). **M)** Histograms summarizing the mean number of mesophyll cells separating veins in WT W22 (blue), *Zmnkd1-Ds;Zmnkd2-Ds* (green), *Zmscr1-m2;Zmscr1h-m1* (yellow) and *Zmscr1-m2;Zmscr1h-m1;Zmnkd1-Ds;Zmnkd2-Ds* (purple) mutants. Error bars are standard error of the mean. Sample sizes (n=) are biological replicates and letters in the top right corner (beneath sample sizes) of each plot indicate statistically different groups (*P*≤0.05, one-way ANOVA and Tukey’s HSD) calculated using the mean number of mesophyll cells in each genotype.

### Evidence for a NKD-mediated maternal effect on leaf patterning in maize

To assess whether ZmNKD1 and ZmNKD2 function redundantly in the context of leaf patterning, we quantified the percentage of fused veins found in plants homozygous for *Zmscr1;Zmscr1h* but with a range of *ZmNKD1* and *ZmNKD2* genotypes (Fig. 8A). These genotypes were generated by self-pollinating plants that had at least one wild-type copy of either *ZmNKD1* or *ZmNKD2*. Surprisingly, some genotypes displayed more fused veins than others. For example, triple *Zmscr1;Zmscr1h;Zmnkd1;ZmNKD2* mutants derived from either *Zmscr1/+;Zmscr1h;Zmnkd1/+;Zmnkd2/+* or *Zmscr1/+;Zmscr1h/+;Zmnkd1;ZmNKD2* parents exhibited a higher number of fused veins than the three quadruple mutants derived from *Zmscr1/+;Zmscr1h;Zmnkd1/+;Zmnkd2/+* or *Zmscr1/+;Zmscr1h;Zmnkd1/+;Zmnkd2* parents (Fig. 8A, Table S1). Given that phenotyping was undertaken on leaf 3, which is initiated and partially patterned in the embryo, we hypothesised that the inconsistencies observed may reflect a maternal effect. Because all parental plants had a wild-type copy of *SCR1*, any inconsistencies must be mediated by NKD, with NKD protein in the parental plant either directly or indirectly compensating for the lack of NKD in the developing embryo.

**Figure 8.**
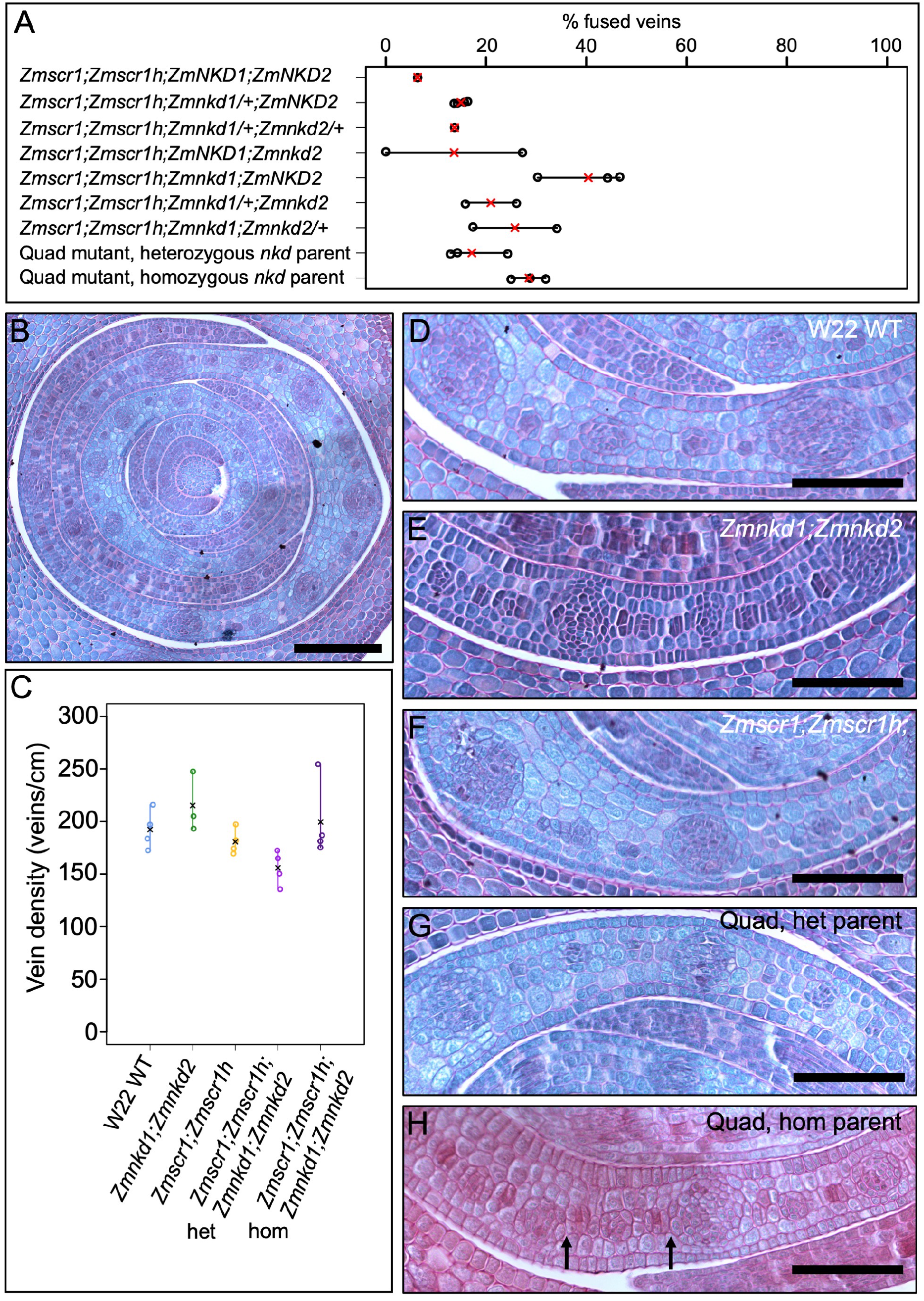
Embryonic leaf phenotypes reveal a NKD-mediated maternal effect. **A)** Quantification of the % of fused veins in leaf 3 of various genotypes. Open circles are data points from distinct biological replicates, red crosses indicate the mean for each genotype. Note that WT have no fused veins (see Fig. 7K). **B)** Transverse section of a wild-type (WT) W22 embryo taken across the tip of the meristem, such that P1-P4 are visible. Scale bar: 100 μm. **C)** Quantification of vein density in P4 primordia. Open circles indicate measurements from independent biological replicates, and black crosses indicate the mean for each genotype. **D-H)** Transverse sections of P4 primordia from WT W22 (D), *Zmnkd1-Ds;Zmnkd2-Ds* (E), *Zmscr1-m2;Zmscr1h-m1* (F), *Zmscr1-m2;Zmscr1h-m1;Zmnkd1-Ds;Zmnkd2-Ds* from a heterozygous *Zmscr1-m2+;Zmscr1h-m1;Zmnkd1-Ds /+;Zmnkd2-Ds/+* parent (G) and *Zmscr1-m2;Zmscr1h-m1;Zmnkd1-Ds;Zmnkd2-Ds* from a homozygous *Zmscr1-m2/+;Zmscr1h-m1;Zmnkd1-Ds;Zmnkd2-Ds* parent (H). Fused veins are noticeable in (H) (black arrows). Scale bars: 50 μm.

To further evaluate the hypothesis that NKD may have a maternal influence on patterning of embryonic leaves, we examined the progeny of a plant which had the genotype *Zmscr1-m2/+;Zmscr1h-m1;Zmnkd1-Ds;Zmnkd2-Ds*. The progeny of this homozygous *nkd1/nkd2* parent segregated ¼ for quadruple mutants and in leaf 3 of each of the three quadruple mutant seedlings examined a greater percentage of fused veins was observed than in leaf 3 of the quadruple mutant seedlings that had originated from a parent with at least one wild-type copy of *NKD1* and/or *NKD2* (Fig. 8A). To further investigate this effect, we compared cross sections of quadruple mutant embryos from a *Zmscr1-m2/+;Zmscr1h;nkd1/+;nkd2/+* (*nkd* heterozygous) parent and a *Zmscr1-m2/+;Zmscr1h;nkd1;nkd2* (*nkd* homozygous) parent, with wild-type, *Zmscr1-m2;Zmscr1h-m1* and *Zmnkd1-Ds;Zmnkd2-Ds* embryos (Fig. 8B-H). Transverse sections from the tip of the meristem (Fig. 8B), where P1-P4 leaf primordia (where P4 will go on to form leaf 1, P3 leaf 2 etc.) are visible, were examined. Due to the anticipated challenges of identifying fused veins in P4 leaf primordia, where cell-division and differentiation is ongoing, we first quantified vein density to test whether it could be used as a proxy for an increase in fused veins. However, no consistent change in vein density was observed between genotypes (Fig. 8C). Sections were therefore carefully examined for evidence of vein fusion by three independent people, two of whom had no knowledge of which sections corresponded to which genotype. All three observers identified fused veins in what were revealed to be the quadruple mutants from a homozygous *nkd* parent, but not in the quadruple mutants from the heterozygous *nkd* parent. Taken together, these results support the hypothesis that there is a maternal effect of NKD on embryonic leaf development. However, progeny of all homozygous *Zmnkd1-Ds;Zmnkd2-Ds* plants develop in the context of a shrunken kernel which has a disordered aleurone comprised of multiple cell layers. As such, the fused vein phenotype in leaf 3 of quadruple mutant progeny could result as an indirect consequence of the altered kernel structure as opposed to a direct consequence of loss of NKD function. To evaluate whether NKD has a direct role, we examined leaf 6 of quadruple mutants that were derived from a homozygous *nkd* parent. These leaves, which are patterned post-germination, also had an increased proportion of fused veins relative to double *Zmscr1;Zmscr1h* mutants (Fig. S8C-F), demonstrating a role for NKD in leaf patterning beyond embryogenesis. Although this observation does not preclude the possibility that the maternal effect is a consequence of physical alterations to the aleurone, collectively these data suggest that NKD functions alongside SCR to pattern both embryonic and post-embryonic leaves in maize.

## DISCUSSION

SCARECROW has recently emerged as a regulator of monocot leaf development (Hughes et al., 2019; Hughes & Langdale, 2022; Slewinski et al., 2012; Wu et al., 2019). Here we have demonstrated that unlike in maize and rice where SCR has distinct functions in inner leaf (maize) or stomatal (rice) patterning, in the C_4_ monocot *Setaria viridis* it undertakes both patterning functions (Fig. 1 & 2). Prior to this study, no regulators that function alongside SCR in monocot leaf patterning had been identified. Here we show that *NKD IDD* genes play such a role, at least in C_4_ monocots (Fig. 3). Although loss of function *nkd* mutants in maize, setaria and rice show no obvious perturbations in leaf development (Fig. 4), when mutations are combined with loss of function *scr* mutations, synergistic interactions are observed in both maize and setaria, but not in rice (Fig. 5-7). Most strikingly, *scr;nkd* mutants in maize and setaria exhibit an increase in the proportion of fused veins, where the bundle-sheath cells of adjacent vascular bundles are in direct contact with no intervening mesophyll cells (Fig. 6 & 7). Intriguingly, the presence or absence of this phenotype in embryonic leaves suggests that the expression of *NKD* in maternal tissue may influence the development of leaves in the embryo (Fig. 8). Taken together, our results provide insight into the evolution of SCR function in monocot leaves and identify another component of the patterning pathway.

We previously proposed that the role of SCR in patterning inner leaf tissues of maize may represent a C_4_-specific function, and that the stomatal role in rice may be a C_3_-specific function (Hughes & Langdale, 2022). The analysis presented here argues against the latter because *Svscr1;Svscr2* mutants of the C_4_ species *Setaria viridis* do not develop stomata. However, inner leaf patterning perturbations were also observed in *Svscr1;Svscr2* mutants, and thus it remains plausible that the inner leaf patterning function is C_4_-specific. In this scenario, the ancestral role of SCR would be in epidermal patterning, with recruitment into Kranz patterning occurring in those monocots that evolved the C_4_ pathway. This situation contrasts with that seen in eudicots where Arabidopsis (C_3_) *scr* mutants develop normal stomata but have perturbations in bundle-sheath cell identity (Cui et al., 2014; Dhondt et al., 2010). On the basis of gene expression patterns and mutant phenotypes in rice, it has been proposed that SCR functions in the epidermis to interpret a positional signal (likely SHR) emanating from veins in the inner leaf (Nunes et al., 2020; Schuler et al., 2018). In this way, stomatal files are correctly positioned in rows adjacent to underlying veins. It is difficult to envisage how this could happen in setaria if SCR is also expressed in the inner leaf as in maize, particularly given that the canonical model for SCR function dictates that SCR prevents SHR movement beyond the cell layer in which SCR accumulates (Cui et al., 2007; Gallagher et al., 2004; Koizumi et al., 2012; Nakajima et al., 2001). How epidermal and inner leaf patterning processes are both mediated through the SCR pathway in setaria is a question for future research.

Another outstanding question from this study is why the severity of the fused vein phenotype is much higher in maize than in setaria *scr;nkd* mutants. In maize, many quadruple mutants exhibited more than 30% of leaf veins that were fused to an adjacent vein, whereas in setaria these incidents were rarer, and no examples were found of more than two veins fused together. One possible explanation is that because *scr;nkd* mutants in setaria have no stomata they are much weaker than those in maize. As a consequence, fewer plants survived and inner leaf phenotypes were necessarily assessed on those survivors, possibly biasing the analysis towards mutants that were less severely affected. An alternative possibility is that the mutant alleles in setaria are hypomorphic rather than null. The successful edits in *SvSCR* and *SvNKD* genes were positioned towards the 3’ end of the sequence, affecting the final 20% and 40% of the encoded proteins respectively. Whereas we can be confident that the *Svscr* alleles are null due to the conserved phenotype of *Svscr1;Svscr2* mutants with rice and maize, it is possible that the *Svnkd* alleles retain some wild-type function and that a more severe *Svscr;Svnkd* vein clustering phenotype would be observed if edits towards the 5’ end of the gene had been successful. If neither of these technical explanations is correct, the differences observed between maize and setaria must represent biological differences between the two species, which is plausible given that they are separated by millions of years of evolution and clearly deploy the SCR pathway in different developmental contexts.

IDD genes have long been known to modulate the SCR/SHR pathway in Arabidopsis. For example, JACKDAW is known to physically interact with the SCR-SHR complex (Welch et al., 2007), and it has been suggested that this interaction with an IDD C2H2 transcription factor partner enables SCR-SHR to bind DNA and activate expression of downstream targets (Hirano et al., 2017). There are IDD orthologs in monocot genomes, but very little functional insight into their roles. This may be in part due to the extensive functional redundancy in grass genomes which combined with the challenges of generating loss-of-function lines in monocots makes functional analysis challenging. However, it was recently found that IDD12 and IDD13 act alongside SHR in rice to regulate ground meristem proliferation in leaves (Liu et al., 2022). This finding is intriguing because we find here that *NKD* plays a similar role alongside *SCR* in C_4_ inner leaf patterning, but that neither gene functions in the inner leaf of rice (Fig. 6, Hughes & Langdale, 2022). It may be that different IDD genes fine-tune the action of both SCR and SHR by controlling the activation of distinct downstream targets in different developmental contexts.

The accumulation of *SCR* and *NKD* transcripts in the ground meristem cells between developing veins, and the fused veins observed in both maize and setaria *scr;nkd* mutants imply two possible mechanisms for SCR and NKD function. The first is that the combined action of SCR and NKD promotes cell division in the ground meristem and then specifies those cells as mesophyll. In this model, in the absence of SCR and NKD ground meristem cells do not divide and/or are mis-specified such that veins develop in direct contact with each other. This would be analogous to the role played by SCR in the arabidopsis root, where loss of gene function leads to loss and mis-specification of the endodermal cell-layer (Laurenzio et al., 1996). The second is that SCR and NKD act to inhibit the formation of veins in regions between existing veins. In their absence, veins are formed in regions between already specified veins, resulting in the runs of fused veins observed in maize and setaria *scr;nkd* mutants. This mechanism may be analogous to the lateral inhibition mechanism that patterns trichomes and stomata on the leaf surface (Schellmann et al., 2002; Torii, 2012; Zeng et al., 2020), where the fused veins represent a clustering phenotype caused by absence of the inhibitor. Because SCR and NKD do not accumulate in developing veins, in this scenario it is likely that they would interpret an inhibitory signal emanating from adjacent specified veins. In either case, the inner leaf perturbations observed in *scr* but not *nkd* mutants suggest that SCR can compensate for loss of NKD function in the inner leaf patterning pathway whereas NKD cannot compensate for loss of SCR function, a relationship that is similar to that seen in other pathways where SCR and IDD proteins interact (Long et al., 2015; Ogasawara et al., 2011; Welch et al., 2007).

NKD has a well-defined role in the endosperm of maize, where it acts to suppress proliferation of the aleurone layer – the aleurone is a single cell layer in wild-type endosperm whereas in *nkd1;nkd2* mutants it is multi-layered (Yi et al., 2015). Within the endosperm, *NKD1* and *NKD2* are expressed in both the aleurone and the starchy endosperm, and loss of function of either gene is compensated for by upregulation of the other (Yi et al., 2015). Although low levels of *NKD* gene expression have also been reported in the developing embryo (Yi et al., 2015), the data reported here suggest that maternally expressed *NKD* may play a role in the patterning of embryonic leaves. Without further experimentation it is impossible to determine the mechanistic basis of this non cell-autonomous effect. Perturbations to leaf patterning in embryos developing within *nkd1;nkd2* mutant seed could either be an indirect consequence of disrupted signalling through the mutant aleurone layer or a direct consequence of loss of activity of NKD and/or its downstream targets. Speculation at this stage would be premature but it is interesting to note that most of the maternal effect genes reported to date encode epigenetic regulators or small molecules (peptides or miRNAs) as opposed to transcription factors (reviewed in Ingram, 2020). Taken together, the data presented here suggest that NKD is a key regulator of leaf patterning in C_4_ grasses, acting with SCR to promote mesophyll cell specification or repress vein specification, and influencing the patterning of embryonic leaves in a non-cell autonomous manner.

## MATERIALS & METHODS

### Phylogenetics

Primary transcript proteomes from *Zea mays* (B73), *Sorghum bicolor*, *Setaria italica*, *Setaria viridis*, *Oryza sativa*, *Brachypodium distachyon*, *Brachypodium stacei*, *Ananas comosus*, *Arabidopsis thaliana*, *Solanum lycopersicum* and *Physcomitrella patens* were downloaded from Phytozome12 (Goodstein et al., 2012). The ZmNKD1 (GRMZM2G129261) primary protein sequence was used as a query in a BLASTp search (e-value of 1e-3) against these proteomes. The top 100 results were aligned using MAFFT-linsi (Katoh & Standley, 2013) and the alignment was then used to generate a maximum likelihood phylogeny using IQtree (Hoang et al., 2018; Trifinopoulos et al., 2016).

### Plant material and growth conditions

Maize inbred line W22 and *Zmnkd1-Ds;Zmnkd2-Ds* (introgressed into W22) seed were obtained from Phil Becraft, Iowa State University. *Zmscr1-m2;Zmscr1h-m1* seed were generated and reported previously (Hughes et al. 2019). W22 was used as the wild-type control in all maize experiments except for *in situ* hybridization experiments where the inbred line B73 was used. *Setaria viridis* accession ME034V was used as the wild-type control for all setaria experiments and *Oryza sativa spp japonica* cv. Kitaake was used as the wild-type control for all rice experiments. *Osscr1;Osscr2* seed were generated and reported previously (Hughes & Langdale, 2022).

Maize and setaria plants for phenotypic analysis were grown in a greenhouse in Oxford, UK with 16 h light/ 8 h dark cycle, daytime temperature 28 °C and night-time temperature 20 °C, with supplemental light provided when natural light levels were below 120 μmol photon m^−2^ s^−1^(Hughes et al., 2019). Maize seed were germinated and grown as described previously (Hughes et al., 2019). Setaria seed were treated for at least 3 weeks in damp sphagnum moss (Zoo-Med Laboratories Inc) at 4°C prior to germination to break dormancy. Seed were then germinated on damp paper towels in sealed petri dishes, in a growth cabinet with the same conditions as the greenhouse. After seven days, seedlings were transferred to Sinclair compost in 60 well modular trays for growth in the greenhouse. Plants grown for seed propagation were re-potted after 4 weeks into 7.5cm pots with the same compost, for growth to maturity. Rice seed were sterilised prior to germination on ½ MS media and subsequently transferred to a hydroponic growth system in 50 ml falcon tubes, as described previously (Hughes & Langdale, 2022).

### Generation of maize quadruple mutant and genotyping

Pollen from *Zmscr1-m2/+;Zmscr1h-m1* plants (double mutants do not produce pollen or ears) was used to cross to *Zmnkd1-Ds;Zmnkd2-Ds* homozygous ears. 50% of the F1 plants were thus heterozygous for all four mutant alleles, and these plants were self-pollinated. In the subsequent F2, kernels with the shrunken *nkd1/nkd2* phenotype were selected for self-pollination. F3 seed derived from individuals with *Zmscr1/+;Zmscr1h;Zmnkd1;Zmnkd2* or *Zmscr1;Zmscr1h/+;Zmnkd1;Zmnkd2* genotypes were then used for quadruple mutant analysis. Other mutant combinations were derived from parental genotypes as indicated in Table S1.

Genomic DNA for genotyping was extracted either using a modified SDS 96-well plate method or a CTAB method depending on the number of samples to be processed, as described previously (Hughes et al 2019). PCR genotyping assays were used to track the genotype at each locus through the generations. The *Zmscr1-m2* and *Zmscr1h-m1* alleles were genotyped as described previously (Hughes et al. 2019). The expected *Ds* insertion site in the *Zmnkd1-Ds* allele was confirmed by two separate PCR assays, one using two primers (nkd1-F, TATCTTATCCGTCGATGCGTTG and nkd1-R, TCGGTCATGGCATCCTGCCTCCG) that flanked the insertion site and produced an amplicon for the wild-type *ZmNKD1* sequence, and another using one flanking primer (nkd1-F) and a second primer nested within the Ds transposon sequence (W22-Ds-R1, GGAGCTGGCCATATTGCAGTCATC) that produced an amplicon when the *Zmnkd1-Ds* sequence was present. The *Zmnkd2-Ds* allele referred to here was previously described as *nkd2-Ds0766* and believed, like *Zmnkd1-Ds*, to be a loss-of-function allele caused by the insertion of a *Ds* transposon. However, attempts to amplify from the *Ds* transposon sequence to the region flanking the insertion failed. Instead, we demonstrated that loss-of-function of this allele was due to a deletion at the 5’ end of the gene, with amplification (primers ZmNKD2-F2, CTTGTTGCCGTTGTTGATTG and ZmNKD2-R2, GTGCCATGTGGCTCCTATTT) yielding a fragment that was 786bp smaller in homozygous *Zmnkd2-Ds* plants than in wild-type (Fig. 4A, Figure S9). This deletion is believed to be caused by a chromosomal rearrangement (Phil Becraft, personal communication). As expected, *Zmnkd1-Ds;Zmnkd2-Ds* seed always exhibited the shrunken kernel phenotype characteristic of *Zmnkd1;Zmnkd2* double mutants. PCR amplifications were undertaken using GoTaq DNA polymerase (Promega) and PCR cycles of 95 °C for 5 min; 35 cycles of 95 °C for 30 s, 57-64 °C for 30 s and 72 °C for 60-90 s; and 72 °C for 5 min. Betaine (1 M; Sigma Aldrich) was added to all reactions to aid amplification of regions with high-GC content.

### Generation of rice and setaria gene edited lines

Rice and setaria *NKD* orthologs were identified from the phylogeny presented in Figure 3. Setaria *SCR* orthologs were identified from a previously published phylogeny (Hughes et al. 2019). The *OsNKD* sequence was obtained from phytozome V12 and two guide RNAs targeting the 5’ end of the *c*oding sequence were designed using CRISPOR (Concordet & Haeussler, 2018) (OsNKD-g59: CTTCGGGATTAGCAGGGATG, OsNKD-g72: GTGATATCAGCGGCTTAATC). Guide RNAs targeting *OsSCR1* and *OsSCR2* were designed and used previously (Hughes & Langdale, 2022), and two of these (OsSCR1-g397: TCCACCCAAGCCGTACTAGG, OsSCR2-g507: CGAGGTCGGGGTTACATGGC) were used in this study. Guides were cloned as described previously into four constructs targeting either *OsNKD* (EC17821: OsNKD-g59 and EC17822: OsNKD-g72) or *OsSCR1, OsSCR2* and *OsNKD* (EC17827: OsSCR1-g397, OsSCR2-g507, OsNKD-g59 and EC17828: OsSCR1-g397, OsSCR2-g507, OsNKD-g72). *SvSCR1, SvSCR2* and *SvNKD* sequences were obtained from phytozome V12, however, as the ME034V accession used for transformation is not the same as the sequenced accession, we first amplified the target regions of the ME034V gene sequences. Surprisingly, we found that the ME034V *SvSCR1* and *SvSCR2* sequences more closely matched the published *Setaria italica* sequences, and the ME034V *SvNKD* sequence was intermediate between the published *viridis* and *italica* sequences. Confirmed ME034V sequences were used in subsequent guide RNA design.

Due to a low editing efficiency of guide RNAs in *Setaria viridis* (Daniela Vlad, personal communication), multiple guides were designed against each gene (Figure S1). Because *SvSCR1* and *SvSCR2* have high sequence similarity, a common set of six guides were designed to target both genes simultaneously. For both *SvSCR* and *SvNKD*, guides were positioned towards the 5’ end of the gene to make complete loss-of-function alleles more likely. However, at least one guide was positioned closer to the 3’ end of the gene both to increase the chance of obtaining mutations and to enable the potential excision of most of the gene sequence. To minimise the number of constructs for transformation, guides were cloned into polycistronic expression modules separated by tRNA spacers (Hahn et al., 2020). Two such polycistronic guide arrays were assembled, one with the six *SvSCR* guides and one with the four *SvNKD* guides (Figure S1), both driven by the rice U3 promoter. These arrays were then combined into three constructs using the same Golden Gate cloning system described previously for rice (Hughes & Langdale, 2022): C402052 (SvSCR array), C402053 (SvNKD array) and C402054 (SvSCR and SvNKD arrays). Three identical constructs were cloned that included the GRF-GIF1 fusion previously shown to enhance rice and wheat transformation efficiency (Debernardi et al., 2020). No enhancement in setaria transformation was observed here, however, one line that was generated from one of these constructs (C402057: SvSCR and SvNKD arrays) was used for subsequent phenotypic analysis after the construct (and thus GRF-GIF1 fusion) was segregated away from the mutations.

Constructs were transformed into Kitaake rice seed using agrobacterium strain EHA105 as described previously (Hughes & Langdale, 2022). Constructs were transformed into setaria ME034V using a modified transformation protocol (Finley et al., 2021). In brief, ME034V seed were dehulled and sterilised with 10% bleach plus 0.1% tween for 3 minutes. Seed were placed on callus induction medium (CIM) (4.3 g/L MS salts, 40 g maltose, 35 mg/L ZnSO_4_.7H2O, 0.6 mg/L CuSO_4_.5H_2_0, 0.5 mg/L kinetin, 2 mg/L 2,4-D 4 g/L Gelzan, pH 5.8) and after 4-5 weeks embryonic callus was moved to fresh medium and gelatinous callus removed. After a further 3 weeks, gelatinous callus was again removed and the remaining callus moved to fresh medium for 1-2 weeks prior to transformation. Constructs were cloned into agrobacterium strain AGL1 and grown in liquid medium to an OD of ~0.6. Agrobacteria were then collected by centrifugation and resuspended in 50ml liquid CIM medium with the addition of 40 μm acetosynringone and 0.02 % synperonic acid. Around 100 pieces of calli were added to this suspension and incubated for 5 minutes with occasional rocking prior to drying and transfer to fresh CIM plates with filter paper placed on top of the medium. Calli were co-cultivated for 3 days at 22°C before transfer to CIM selective plates (CIM containing 150 mg/L timentin and 40mg/L hygromycin) for 16 days at 24°C in the dark. Calli were then transferred to plant regeneration medium (PRM) (4.3 g/L MS salts, 20 g/L sucrose, 7 g/L Phytoblend, 2 mg/L kinetin, 150 mg/L timentin, 15 mg/L hygromycin, pH 5.8) and moved to fresh medium every 2 weeks. Emerging shoots were dissected from calli and moved to rooting medium (RM) (2.15 g/L MS salts, 30 g/L sucrose, 7 g/L Phytoblend, 150 mg/L timentin, 20 mg/L hygromycin, pH 5.7). Shoots that survived this stage were transferred to compost for genotyping.

Genomic DNA for rice and setaria T0 genotyping was obtained using the same methods described for maize. T0 seedlings were first screened using primers that amplified a fragment of the hygromycin gene (Hughes & Langdale, 2022) to assess transformation success. Gene specific primers were then used to amplify and sequence the region targeted for editing. In rice, all guides induced successful edits, however, in the case of construct EC17822 only a few transformed plants were obtained and thus edited plants from EC17821 were taken forward for *Osnkd* single mutant characterization. In setaria, only one guide for both *SCR* and *NKD* (SvSCR-ex2g49: GAGCAGGACCTGAGCCACTC and SvNKD-ex3g438: CATGAGTCCATGGAACGGCT) was found to consistently yield successful edits. In both cases this guide was positioned closer to the 3’ end of the gene. In the case of *SCR*, out-of-frame mutations created by the successful guide resulted in the final ~20% of the protein sequence being either nonsense or prematurely truncated by an early stop codon, whereas for *NKD* the final ~40% of the protein was affected. As was the case when generating rice *scr* mutants (Hughes & Langdale, 2022), no plants were identified in the T0 generation that had out-of-frame mutations in all four *SCR* alleles, with all screened plants having at least one unedited copy of one of the *SCR* genes. Interestingly, some T0 plants exhibited pale sectors in leaves, but only in plants transformed with the *SCR* guide array. All *Svnkd* mutations corresponded to the same C deletion (*Svnkd-m1*), and plants homozygous for this edit were viable. For both rice and setaria, T1 lines were screened to identify mutated plants that had segregated away from the construct. From this analysis, two independent *Osnkd* lines, three *Osscr1;Osscr2;Osnkd* lines (alongside two previously generated *Osscr1;Osscr2* lines), two *Svscr1;Svscr2* lines, two *Svnkd* lines and three *Svscr1;Svscr2;Svnkd* lines were prioritized for phenotypic characterization in the T2 and T3 generation.

### Leaf phenotyping

Fully expanded leaves were used to obtain transverse leaf sections of maize, setaria and rice. For maize and setaria a region from the midpoint along the proximal-distal axis was cut and placed in molten 7% agar. Once solidified, blocks were trimmed down and mounted using superglue for sectioning on a vibratome. Sections of 30-60 μm were cleared in either 3:1 ethanol: acetic acid (maize) or 70% ethanol (setaria) for 10 minutes. *Svscr1;Svscr2* and *Svscr1;Svscr2;Svnkd* leaves were sufficiently pale for leaf anatomy to be visible without any clearing. Sections were transferred to slides and imaged using a Leica DMRB microscope with a DFC7000T camera under either brightfield or UV illumination using Leica LASX image analysis software. Rice leaf transverse sections were obtained by first fixing a segment from the midpoint along the proximal-distal axis of fully expanded leaf 5 in 3:1 ethanol: acetic acid for 30 mins, before transferring to 70% ethanol. Samples were wax infiltrated, sectioned at 10 μm using a Leica RM2135 rotary microtome and stained with Safranin O and Fastgreen as described previously (Hughes & Langdale, 2022). Stomatal impressions of setaria leaves were obtained and imaged under phase-contrast microscopy using the same dental resin and nail varnish method described previously (Hughes & Langdale, 2022).

### Seed phenotyping

Setaria seed were and dehulled fixed in FAA (10% formaldehyde, 5% acetic acid, 50% ethanol) for 2 months prior to sectioning. Seed were then embedded in paraffin as described previously (Hughes & Langdale, 2022) and sectioned into 20μm sections using a Leica RM2135 rotary microtome. The resultant sections were incubated in Histoclear (2 x 10 min), and then re-hydrated through 100%, 95%, 90%, 80%, 70%, 50%, 30% and 10% (w/v) ethanol, all at room temperature. Sections were then rinsed in distilled H_2_O, stained for 5 seconds in 0.05% (w/v) Toluidine Blue (50 mM citrate buffer, pH 4.4), and then rinsed in distilled H_2_O again, before finally being mounted using a drop of entellen (Merck Millipore). Slides were imaged under brightfield using the same microscope described above. Rice and setaria whole seed were photographed using a Leica S9i stereo microscope.

### In situ hybridization

*In situ* hybridization was carried out using wax-embedded shoot apices as described by Schuler et al., (2018), with digoxygenin (DIG)-labelled RNA probes designed to specifically detect either *ZmSCR1* or *ZmNKD1* transcripts. The *ZmSCR1* probe was a 108 bp region towards the end of the first exon, which shared 78% identity with the corresponding region of *ZmSCR1h* (Hughes et al., 2019). The *ZmNKD1* probe was a 116 bp region of the 5’UTR ending 12 nucleotides upstream of the ATG. The probe shared 59% identity with the corresponding region of *ZmNKD2* and with the stringency conditions used was expected to be specific for *ZmNKD1*. Post-hybridization washes were undertaken with 0.005 x SSC buffer made from a 20x SSC stock (3M NaCl, 0.3M Na3citrate), calculated to ensure stringency.

### Maize kernel genotyping and embryo imaging

The endosperm of kernels from segregating seed packets was first chipped and used for genotyping using the CTAB DNA extraction method described above, except samples were homogenized prior to the addition of extraction buffer. The resultant genomic DNA was used as template in PCR amplifications to identify homozygous mutants for phenotypic characterization. Kernels of interest were then soaked overnight in water, to enable the removal of the embryo the following day. Embryos were fixed in FAA (10% formaldehyde, 5% acetic acid, 50% ethanol) and vacuum infiltrated before incubation overnight in fresh FAA. Samples were transferred to 50% and then 70% ethanol the next day, and wax infiltrated as described previously (Hughes & Langdale, 2022). 10 μm transverse sections were obtained using a microtome just beneath the tip of the meristem where all primordia are visible, and stained with Safranin O and Fastgreen as described above. Slides were imaged using brightfield illumination and the whole region of interest obtained using the Leica XY builder software.

### qPCR

RNA was extracted using the RNeasy kit (Qiagen) from whole maize shoots of both *Zmscr1-m2;Zmscr1-m1* and *Zmnkd1-Ds;Zmnkd2-Ds* mutants and the respective wild-type lines, 7 days after germination. RNA was DNase treated (TURBO DNase, Thermo Fisher) and used as a template to generate cDNA (Maxima first strand cDNA synthesis kit, Thermo Fisher). Primers were designed to amplify short fragments of *ZmSCR1, ZmSCR1h, ZmNKD1* and *ZmNKD2* gene sequences as well as two housekeeping genes (*ZmCYP* and *ZmEF1α*) verified previously as suitable for use in maize qRT-PCR (Fig. S3A) (Lin et al., 2014). RT-PCR was used to verify that a single product of the correct size was amplified. Quantitative RT-PCR amplifications were undertaken using SYBR-Green (Thermo Fisher) with cycling conditions 95 °C for 10 min, then 40 cycles of 95 °C for 15 s and 60 °C for 1 min. Melt curves were obtained by heating the resultant product from 60 °C to 95 °C to confirm that a single product was amplified for each primer pair (Fig. S3B). Amplification efficiencies were assessed using a dilution series of cDNA template with >80% taken to be suitable. In all cases three technical replicates with Ct range <0.5 were obtained from each sample and all comparisons were run on the same 96-well plate. Ct values were obtained using the qPCR miner algorithm (Zhao & Fernald, 2005) and fold-change between wild-type and mutants was calculated using the 2^−ΔΔCT^ method (Livak & Schmittgen, 2001). A combined wild-type average was used to compare each individual wild-type to indicate the range of the wild-type data. Mutant samples were then compared with the same overall wild-type average. Values <1 indicate a relative reduction compared with wild-type whereas values >1 indicate a relative increase compared to wild-type.

## AUTHOR CONTRUBUTIONS

T.E.H and J.A.L conceived and designed the experiments. O.S carried out the *in situ* hybridization experiment and M.T. analyzed the embryonic leaf phenotypes in maize and the aleurone phenotype in setaria. T.E.H carried out all of the remaining experiments and analysed the data. T.E.H and J.A.L wrote the manuscript.

## ACKNOWLEDGEMENTS

The authors thank Phil Becraft, Erik Vollbrecht, Hao Wu, Ruaridh Sawers and Ruben Rellan Alvarez for enabling maize genetics in the field in Iowa and Mexico; John Baker for plant photography; Roxaana Clayton, Julie Bull and Lizzie Jamison for technical support; Matthew Karadzas for initiating the qRT-PCR experiments; Sophie Johnson, Chiara Perico, Daniela Vlad, Sovanna Tan, Julia Lambret-Frotte and Maricris Zaidem for discussion throughout the experimental work and during manuscript preparation, and Mike Scanlan for helpful comments on the manuscript.

---

**Figure S1.**
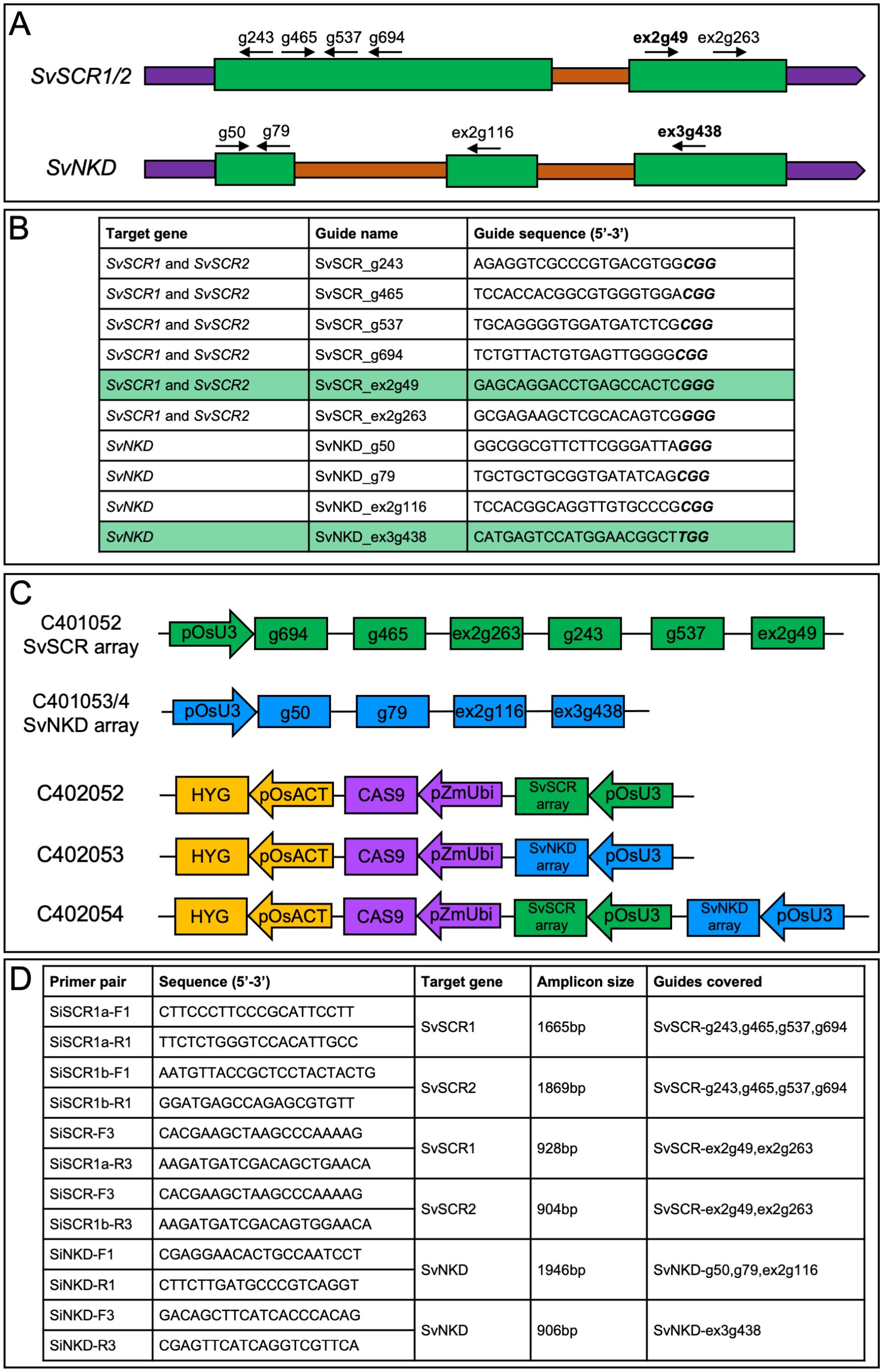
Summary of setaria CRISPR/Cas9 design. **A)** Cartoon summaries of *SvSCR1/2* and *SvNKD* genes with guide positions indicated by black arrows above each gene sequence. 5’ and 3’ untranslated regions are depicted in purple, coding sequences in green and introns in orange. **B)** Guide RNA names and sequences. Guides that successfully edited are highlighted green. **C)** Cartoon summary of Level 1 and Level 2 Golden Gate constructs designed for this study. Promoters are indicated by arrows, and expression modules by rectangles. **D)** Genotyping primers used in this study.

**Figure S2.**
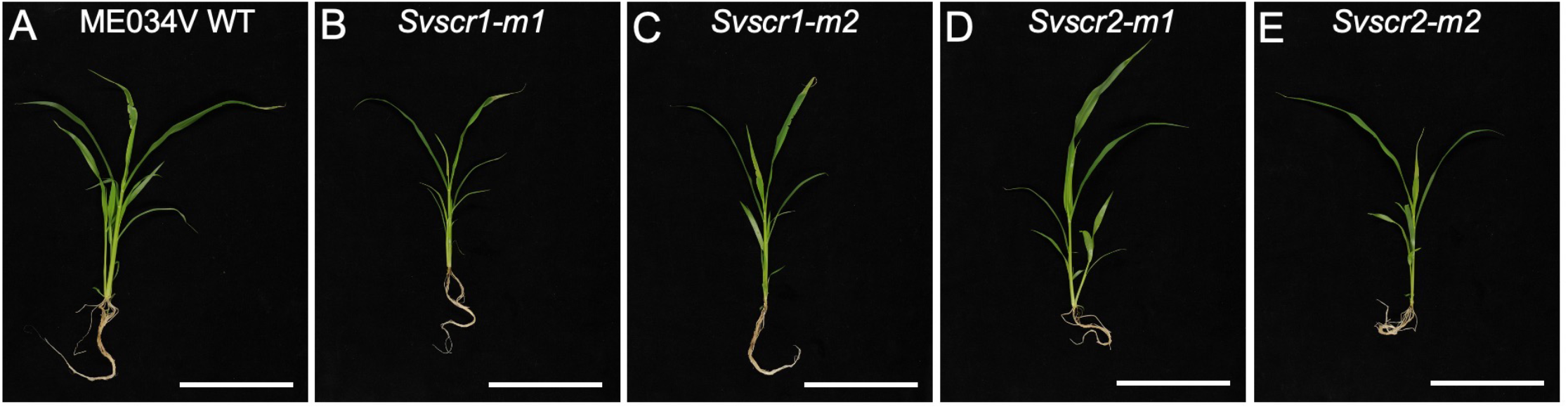
*Svscr1* and *Svscr2* single mutants are phenotypically indistinguishable from wild-type. **A-E)** Images of wild-type (WT) ME034V (A), *Svscr1-m1* (B), *Svscr1-m2* (C), *Svscr2-m1* (D) and *Svscr2-m2* (E) plants taken 21 days after sowing. Scale bars: 10cm.

**Figure S3.**
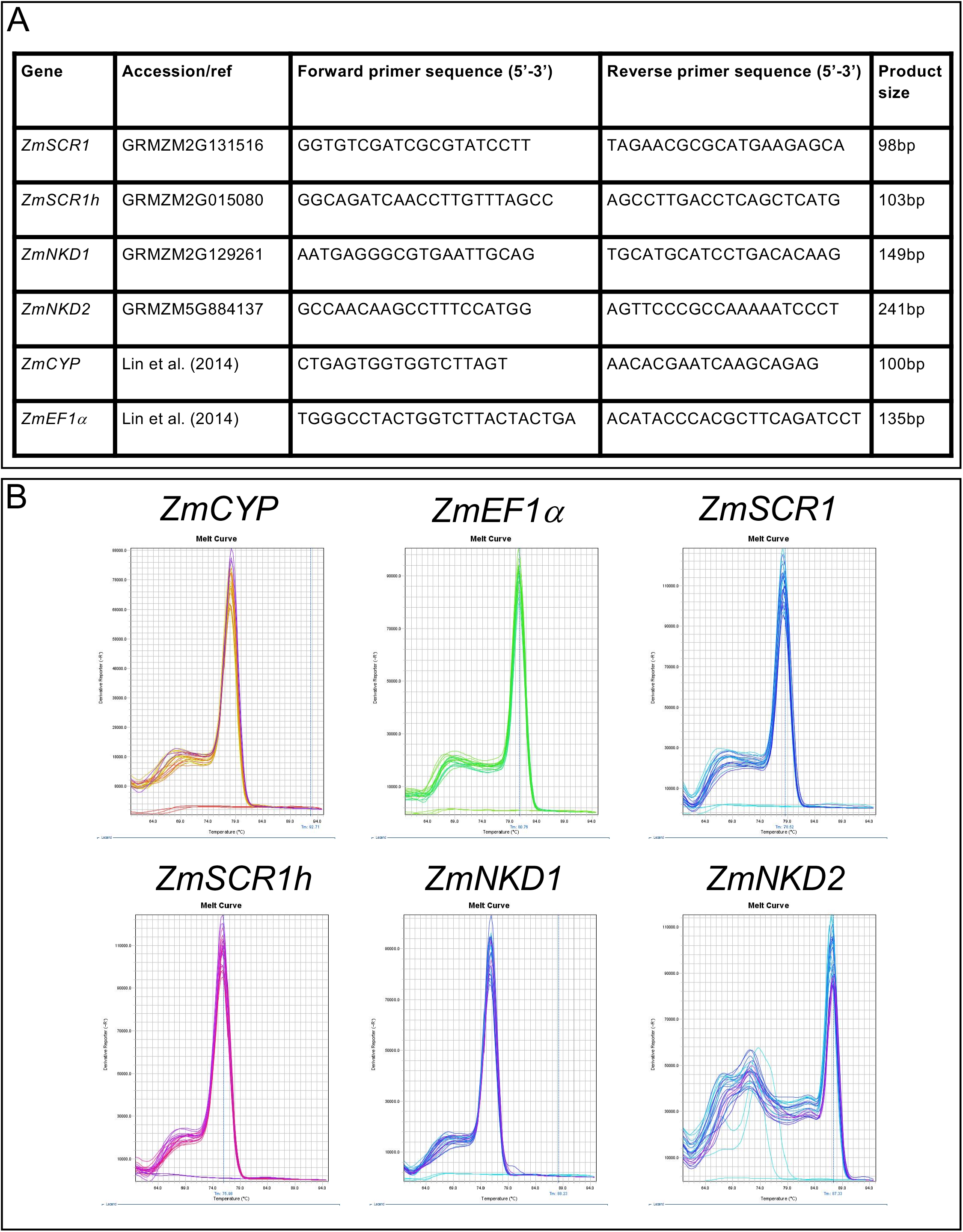
Primers used for quantitative RT-PCR. **A)** Primer sequences used for quantitative RT-PCR of each gene in this study. **B)** Melt-curves for each primer pair.

**Figure S4.**
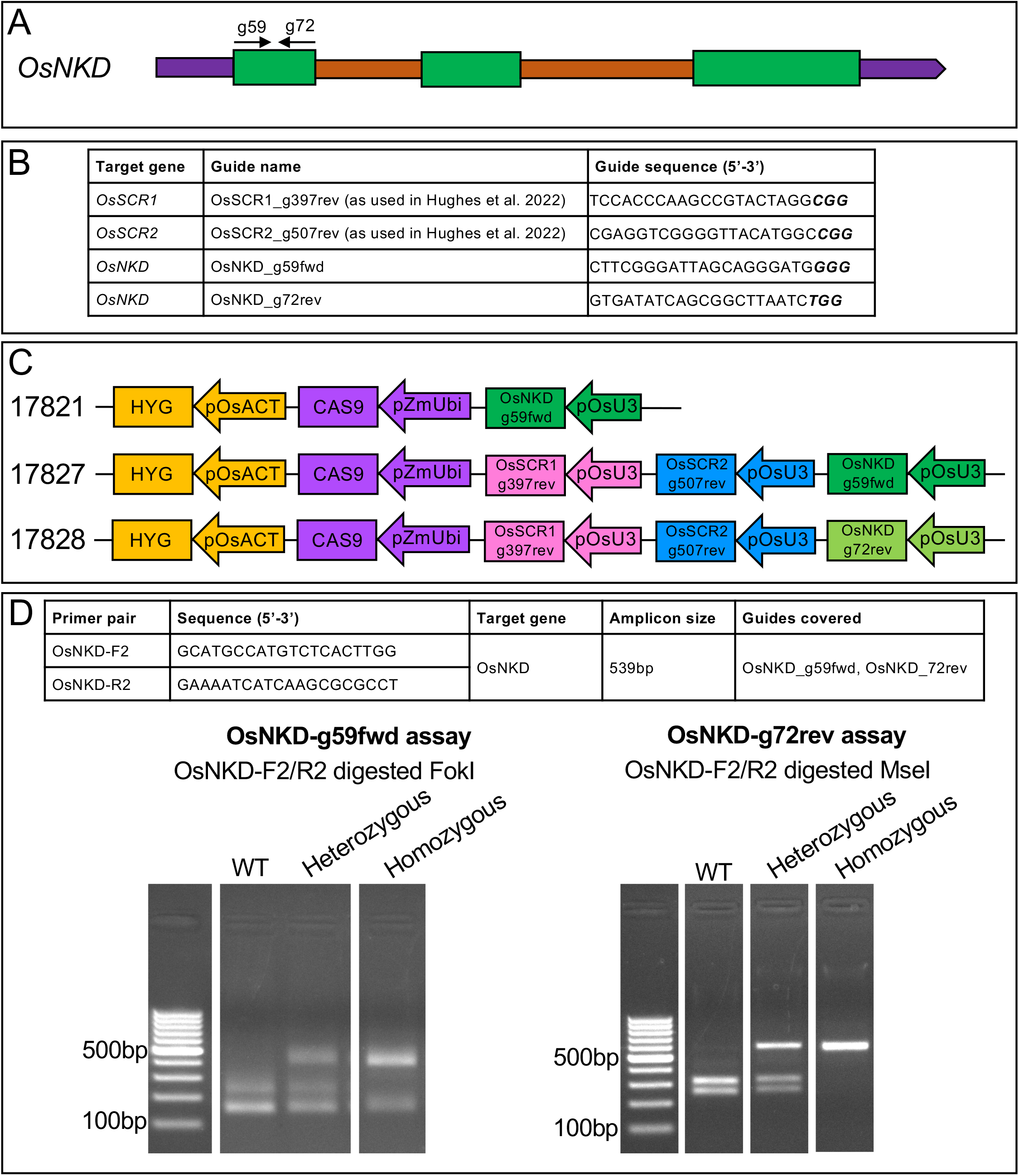
Summary of rice CRISPR/Cas9 design. **A)** Cartoon depiction of *OsNKD* with guide positions indicated above the gene with black arrows. 5’ and 3’ untranslated regions are depicted in purple, coding sequences in green and introns in orange. **B)** Guide RNA sequences used in this study. **C)** Cartoon summary of Level 2 Golden Gate constructs used to generate *Osnkd* (17821) and *Osscr1;Osscr2;Osnkd* (17827 and 17828) mutants. **D)** Primer sequences and restriction digests used to genotype *OsNKD* editing events.

**Figure S5.**
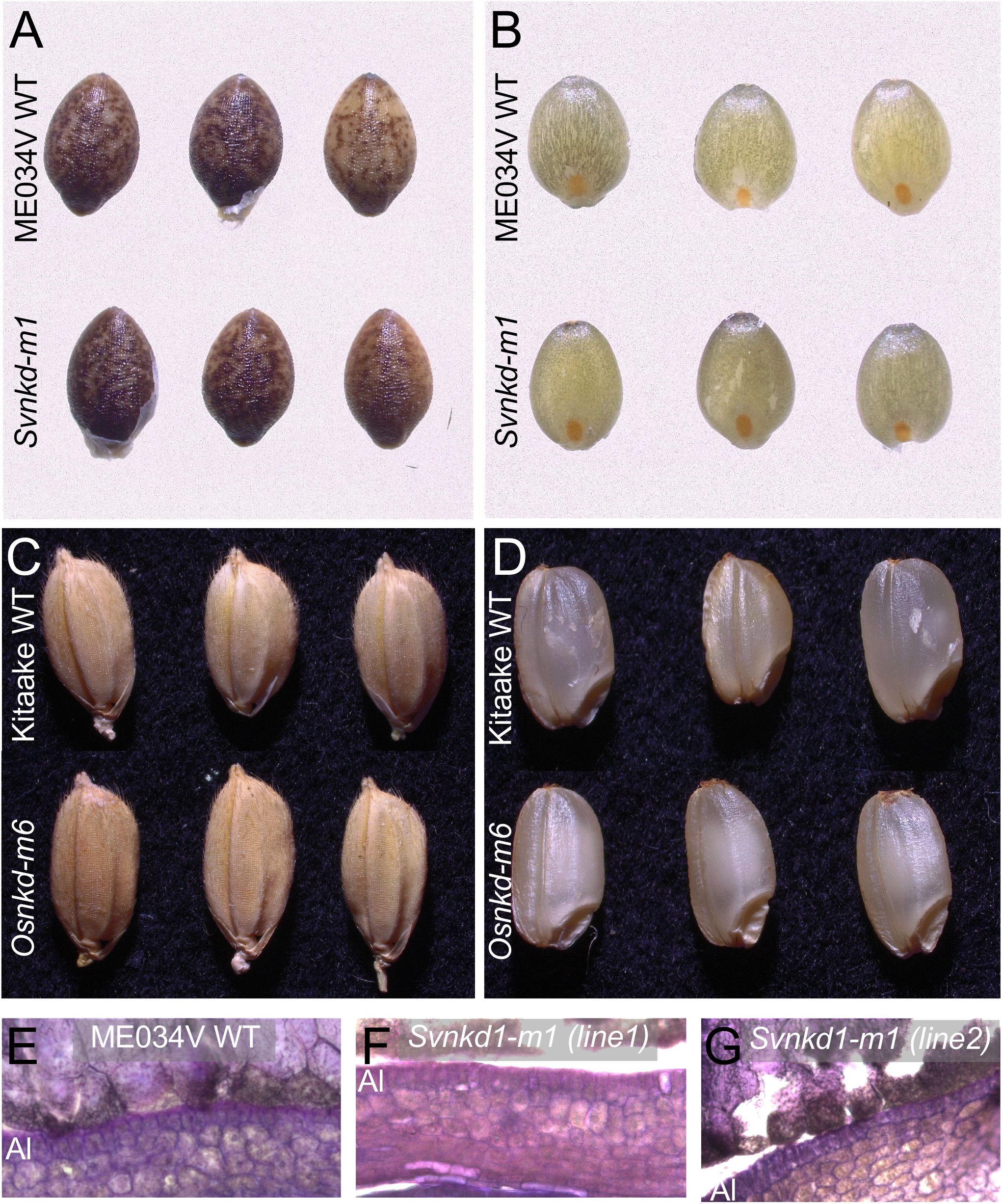
*nkd* mutants do not have perturbed seed development in setaria or rice. **A-D)** Images of setaria (A & B) and rice (C & D) seed with (A & C) or without (B & D) the husk for both wild-type (WT) (top rows) and *nkd* mutants (bottom rows). **E-G)** Cross sections of setaria WT ME034V (E), *Svnkd-m1* (line 1) (F) and *Svnkd-m1* (line 2) (G) mature seed. The aleurone layer is indicated by Al in each image.

**Figure S6.**
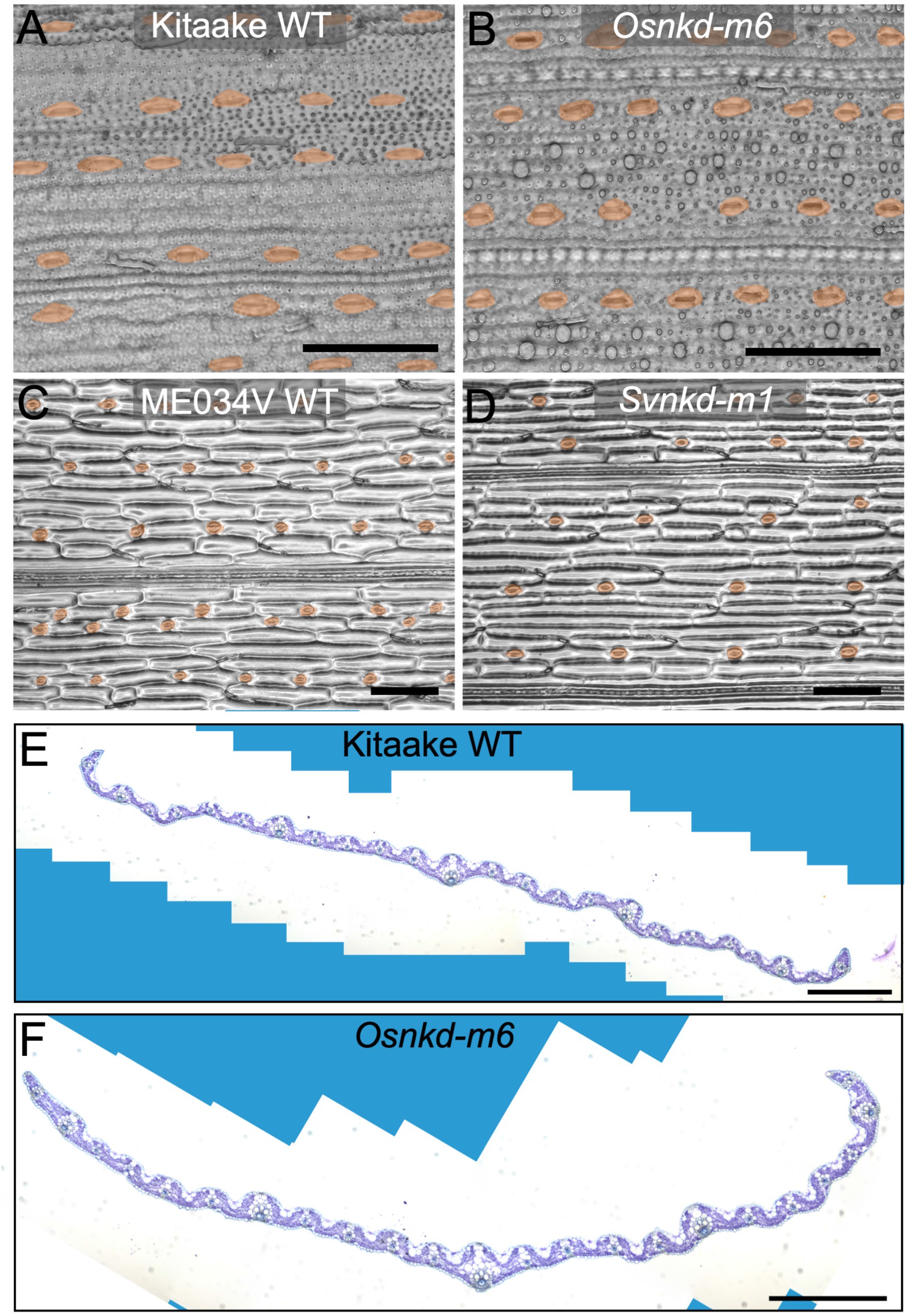
*Osnkd* and *Svnkd* mutants do not exhibit perturbed leaf development. **A-D)** Stomatal impressions of the abaxial surface of wild-type (WT) Kitaake rice (A), *Osnkd-m6* (B), WT setaria ME034V (C) and *Svnkd-m1* (D) leaves. Rice images are taken of leaf 5 and setaria images of leaf 3. Stomata are false coloured orange. Scale bars: 100 μm. **E, F)** Cross sections of WT Kitaake (E) and *Osnkd-m6* (F) leaf 5 taken from the midpoint along the proximal-distal axis. Scale bars: 200 μm.

**Figure S7.**
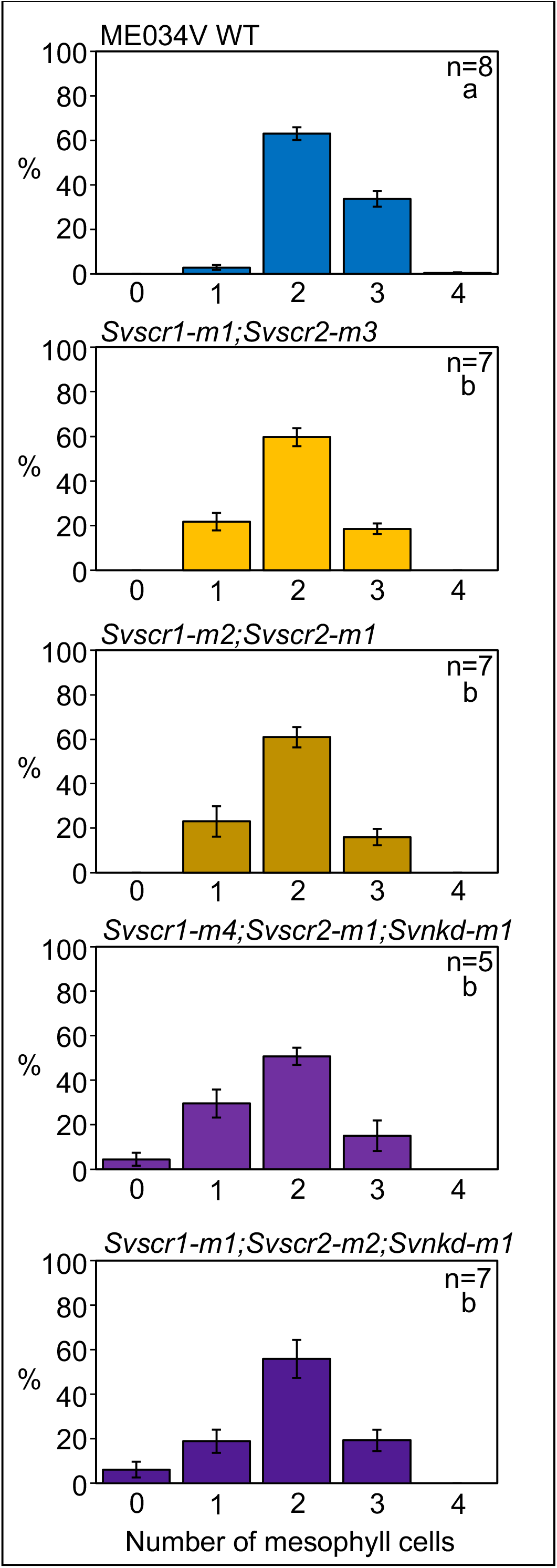
*Svscr1;Svscr2;Svnkd* mutant leaves exhibit occasional fused veins with no intervening mesophyll cells. Quantification of the number of M cells in five setaria genotypes: wild-type ME034V, *Svscr1-m1;Svscr2-m3*, *Svscr1-m2;Svscr2-m1*, *Svscr1-m4;Svscr2-m1;Svnkd-m1* and *Svscr1-m1;Svscr2-m2;Svnkd-m1*. Quantification undertaken on leaf 4. Bars are the standard error of the mean. Biological replicates (n=) are indicated and letters in the top right corner of each plot indicate statistically different groups (*P*≤0.05, one-way ANOVA and Tukey’s HSD) calculated using the mean number of M cells in each genotype.

**Figure S8.**
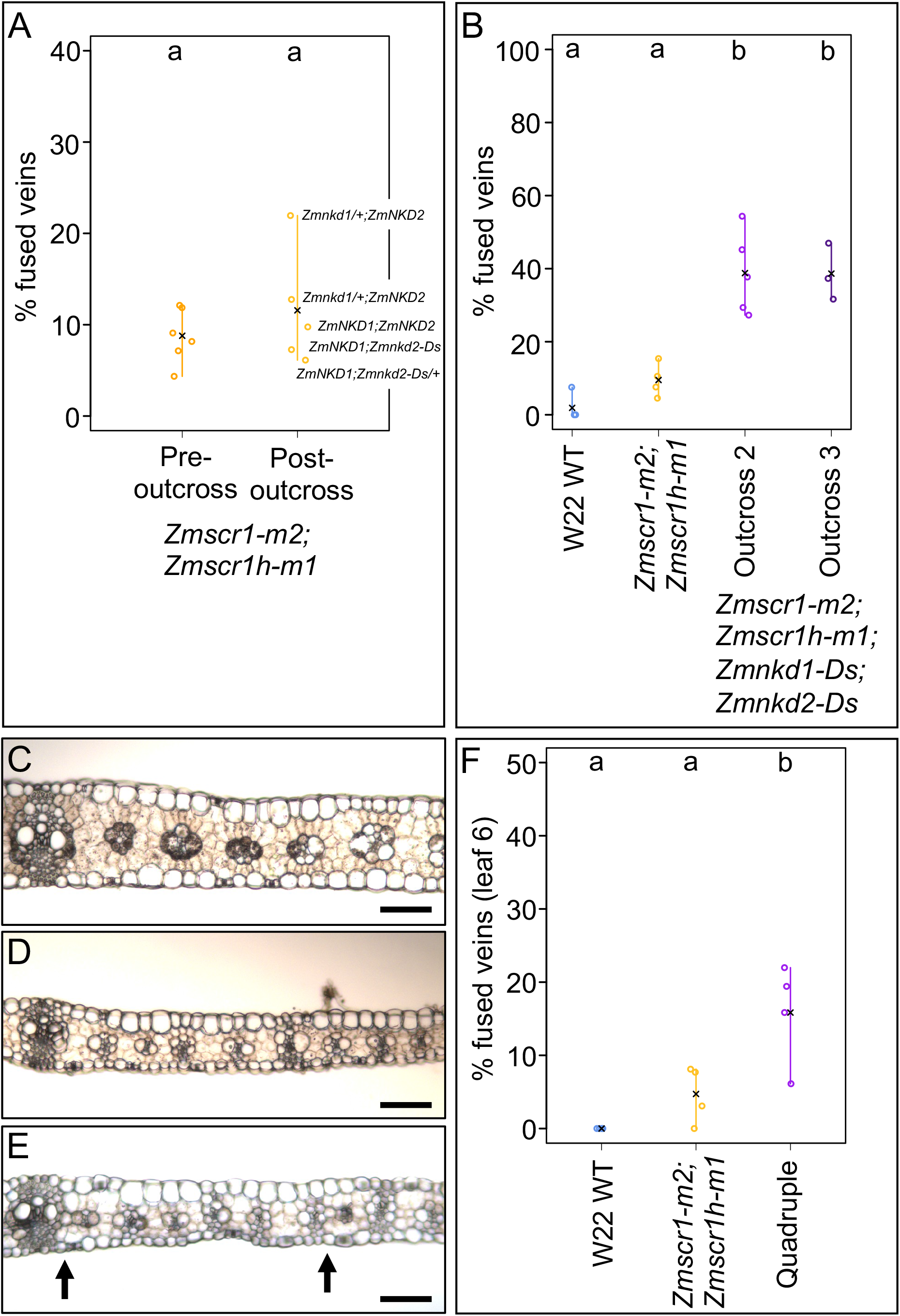
Quadruple *Zmscr1-m2;Zmscr1h-m1;Zmnkd1-Ds;Zmnkd2-Ds* mutants exhibit increased fused leaf veins in a variety of contexts. **A)** Quantification of the percentage of fused veins in leaf 3 of *Zmscr1-m2;Zmscr1h-m1* mutants pre- and post-outcross to *Zmnkd1-Ds;Zmnkd2-Ds*. Mutants phenotyped post-outcross were also segregating for *Zmnkd1-Ds* and *Zmnkd2-Ds* (genotypes labelled on figure). **B)** Quantification of the percentage of fused veins in leaf 3 of wild-type (WT) W22, *Zmscr1-m2;Zmscr1h-m1* and two additional quadruple mutants from two outcrosses using independent *Zmnkd1-Ds;Zmnkd2-Ds* plants. **C-E)** Transverse sections of leaf 6 from WT W22, *Zmscr1-m2;Zmscr1h-m1* and *Zmscr1-m2;Zmscr1h-m1;Zmnkd1-Ds;Zmnkd2-Ds* lines. In (E) fused veins are indicated by arrows. Scale bars: 100 μm. **F)** Quantification of the percentage of fused veins in WT W22, *Zmscr1-m2;Zmscr1h-m1* and *Zmscr1-m2;Zmscr1h-m1;Zmnkd1-Ds;Zmnkd2-Ds* leaf 6. In (A), (B) and (F) open circles are data points from biological replicates, and black crosses indicate the mean for each genotype. Letters above each genotype indicate statistically different groups (*P*≤0.05, one-way ANOVA and Tukey’s HSD).

**Figure S9.**
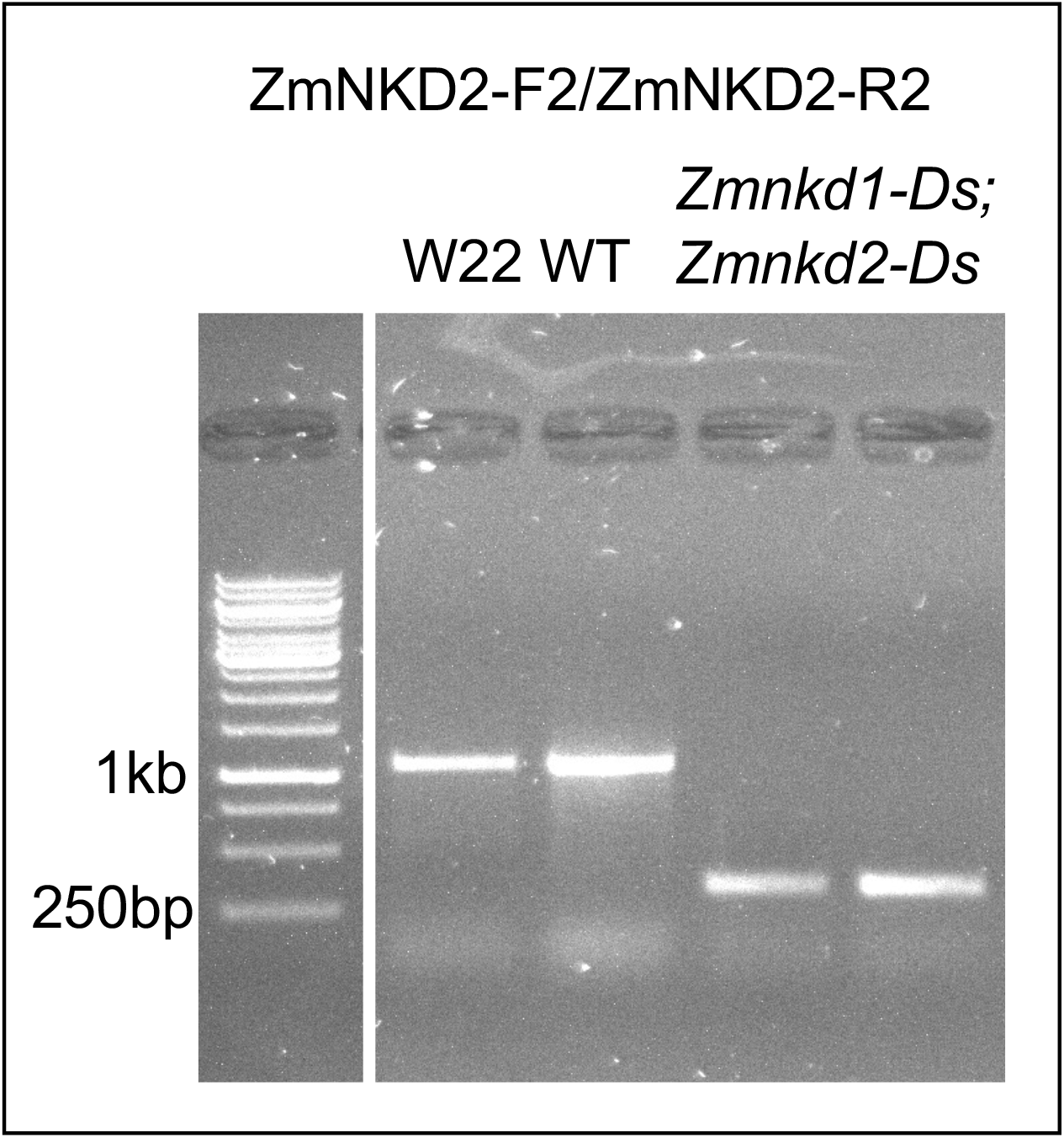
*Zmnkd2-Ds* genotyping. Genotyping of the *Zmnkd2-Ds* allele revealing a 786bp deletion.

**Table S1.**
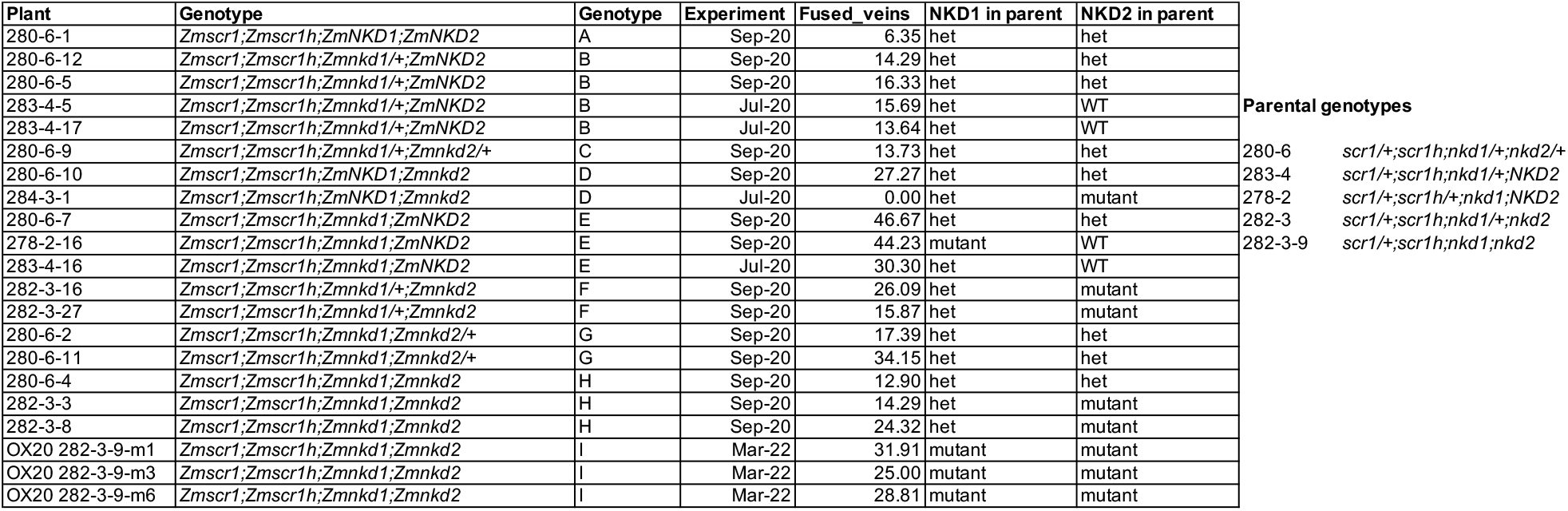
Parental and zygotic genotypes of different combinations of scr and nkd mutant alleles in maize, along with the presence or absence of fused vein phenotypes.

